# Paralogs with distinct phase behaviors broaden the *in vivo* stress response range of condensates

**DOI:** 10.1101/2025.10.13.682129

**Authors:** Jian Guan, Bikash R. Sahoo, Tyler S. Brant, Pavithra Mahadevan, Antonia Brooks, Shyamal Mosalaganti, James C.A. Bardwell, Ursula Jakob

## Abstract

Paralogs are widespread, but their physiological roles are often masked by redundancy. H-NS, a nucleoid-associated protein in Gram-negative bacteria, typically coexists with paralogs such as StpA, whose functions have remained obscured due to the lack of strong phenotypes. We demonstrate here that the interaction between H-NS and StpA fine-tunes the physico-chemical properties of heterochromatin-forming condensates. H-NS forms dynamic condensates, while StpA assembles into stable insoluble fibrils. Yet together, the two proteins form heterotypic condensates, whose fluidity and stability are tunable by their relative stoichiometry. By increasing the levels of StpA over H-NS, bacteria increase heterochromatin-associated condensate stability, thus preserving gene repression and optimizing bacterial growth under stress conditions. Structural differences at these protein’s dimerization sites help explain their distinct phase behaviors. Our findings reveal a novel paradigm in which paralogs that are positioned at the opposite ends of the phase spectrum can fine-tune condensates to promote survival in fluctuating environments.

## INTRODUCTION

Homologous genes within a species, commonly known as paralogs, are pervasive among both prokaryotic and eukaryotic organisms^1,2^. For instance, >60% of human genes and >30% of *Escherichia coli* genes have at least one paralogous sequence within their genomes^3,4^. Acquisition of paralogs, through gene duplication or horizontal gene transfer, is thought to represent an evolutionary response to environmental pressure^1,5,6^ and has also important implications for the virulence and antibiotic resistance of pathogenic bacteria^7–9^. Yet, the functional delineation of members within a paralogous family is often obscured by the lack of phenotypes upon the loss of the individual paralogs. As a result, paralogs are regularly overlooked as functionally redundant proteins, potentially dismissing their contributions to organismal fitness under non-optimal growth conditions^10^.

The Histone-like Nucleoid Structuring protein (H-NS) is an archetypal nucleoid-associated protein (NAP), widely present in Gram-negative bacteria. It consists of an N-terminal oligomerization domain with two distinct dimerization sites (1 and 2) that is linked to a C-terminal DNA binding domain (DBD) via a short intrinsically disordered region (IDR) (Figs. S1A, B). H-NS functions primarily as a transcriptional silencer by oligomerizing into extended filaments along regions of the nucleoid that are enriched in horizontally acquired genes^11–14^. Accordingly, deletion of the *hns* gene leads to a widespread transcriptional derepression^15–17^ and has profoundly negative impacts on bacterial growth, metabolism and virulence^18–20^. In addition to H-NS, *E. coli* and many other enteric bacteria encode the *S*uppressor of mutant *t*d *p*henotype A (StpA) protein, a paralog that shares 58% sequence identity with H-NS (Figs. S1A, C)^21^. StpA and H-NS display highly overlapping DNA binding landscapes^22,23^, consistent with their ability to form heterotypic nucleoprotein filaments^24,25^. Compared to filaments comprised solely of H-NS, mixed H-NS/StpA filaments are more thermostable and show an increased capacity to stall RNA polymerase, likely due to StpA’s higher thermal stability, DNA bridging ability and DNA binding affinity as compared to H-NS^26–28^. Nonetheless, under non-stress conditions, StpA levels are kept low through the repressive function of H-NS as well as its own negative autoregulation^24,29–31^. Moreover, loss of StpA causes negligible transcriptional changes, growth defects or virulence phenotypes^32–35^. For these reasons, StpA has long been considered to serve primarily as a molecular backup of H-NS^36^. However, this simple idea overlooks the poor ability of StpA to compensate for the loss of H-NS^37^. It also fails to explain why StpA is targeted by Lon-mediated proteolysis in the absence of H-NS, suggesting that its interactions with H-NS are necessary to stabilize StpA^38,39^. Interestingly, in response to high temperature or hyperosmolarity, StpA abundance increases dramatically^24,40^, and its deletion begins to impair bacterial fitness^41^. These results made us wonder whether StpA was evolutionarily selected for physicochemical properties that confer H-NS-like functions under non-optimal growth conditions, accepting that these alterations might pose adverse effects and hence require tight control of StpA protein levels under standard growth conditions.

Here, we provide *in vitro* and *in vivo* evidence that H-NS and StpA differ drastically in their phase behaviors. Whereas H-NS on its own forms short linear oligomers that readily assemble into highly fluidic condensates, StpA precipitates in form of long and insoluble fibrils. However, when incubated together, H-NS and StpA associate into short soluble fibrils that form co-condensates at a significantly reduced saturation concentration, and with significantly increased stability towards elevated temperatures and osmolarity than H-NS condensates. We demonstrate that exposure of *E. coli* to heat or osmotic stress increases the StpA to H-NS ratio within the condensates, preserving their gene silencing function under stress conditions that would otherwise destabilize and dissolve H-NS condensates. Molecular dynamic (MD) simulations reveal significant structural differences between the paralogs at dimerization site 2, which explain their different phase behaviors and stabilities. These results provide evidence for organisms being able adjust the stoichiometry of interacting paralogs with distinct phase behavior to expand the functionality of condensates over a wide range of environmental conditions.

## RESULTS

### H-NS and StpA form *in vivo* co-condensates

To simultaneously monitor the *in vivo* behavior of both *E. coli* H-NS and its paralog StpA, we constructed a derivative of the wild type (WT) *E. coli* strain MG1655, which encodes both H-NS tagged at the C-terminus with the monomeric fluorescent protein mNeonGreen (H-NS-mNG) and StpA tagged with mCherry (StpA-mCh) integrated at their respective endogenous loci. We grew this strain, which showed WT-like growth behavior (Fig. S1D) in defined minimal medium, and conducted live cell imaging. As expected for NAPs that interact *in vivo*, their fluorescent signals were strongly associated with each other as well as with the nucleoid (Fig. 1A, upper panels; Fig. S1E). However, we noted that in exponentially growing bacteria, the H-NS-mNG and StpA-mCh signals were not evenly distributed along the nucleoid but were enriched in submicron domains and in foci that appeared to undergo constant dynamic structural rearrangements (Fig. S1F). Fluorescence recovery after photobleaching (FRAP) experiments on the portions of the nucleoid that contained enriched H-NS-mNG and StpA-mCh fluorescent signals revealed significant fluorescence recovery, confirming that both proteins form dynamically exchangeable sub-compartments, suggestive of *in vivo* condensates (Figs. S1G, H). These results were in line with recent reports, which showed that purified H-NS phase-separates and forms reversible condensates with DNA *in vitro*^42^. Live cell imaging of the same strain in late stationary phase (i.e., 24h after OD_600_ = 0.2) showed that under these conditions, H-NS-mNG and StpA-mCh form even more highly distinct and colocalized puncta that align with the longitudinal axis of the bacteria (Fig. 1A, lower panel). Both NAPs were predominantly soluble (Fig. S1I), making it unlikely that these puncta represent protein aggregates. Moreover, we observed distinct fusion and fission events among these puncta, albeit happening more rarely and slower than those observed during exponential phase (Fig. S1J). FRAP experiments further confirmed the fluidic nature of these condensates by showing the recovery of both fluorescent signals (Fig. 1B, C). Of note, however, we observed that the H-NS-mNG fluorescence recovered to a significantly higher extent than the StpA-mCh signal, suggestive of a potentially paralog-specific difference in their dynamic diffusivities. Upon recovery from stationary phase, both H-NS and StpA signals simultaneously dispersed from the puncta, and, within 4h of recovery, revealed the more diffuse pattern resembling that observed during exponential growth (Fig. 1D). Distinct H-NS/StpA foci also formed upon cultivation of the strain in rich media (Fig. S1K) and were detected in fixed bacteria that express H-NS fused to a much shorter 3xFLAG tag (Fig. S1L), making it unlikely that the observed puncta are caused by the fluorescent protein tags. Furthermore, stationary-phase *E. coli* strains expressing H-NS-mNG in concert with mCherry-tagged fusions of other known oligomerization-competent *E. coli* NAPs demonstrated that these unrelated fusion proteins either did not form distinct foci (i.e., HupA- mCh, IhfA-mCh, Lrp-mCh, Fis-mCh) or formed puncta that did not colocalize with H-NS-mNG (i.e., Dps-mCh, Hfq-mCh) (Fig. S1M). Collectively, these data provide evidence that the paralogous nucleoid-associated proteins H-NS and StpA engage in the formation of heterotypic condensate-like assemblies with potentially distinct dynamic properties.

**Figure 1.**
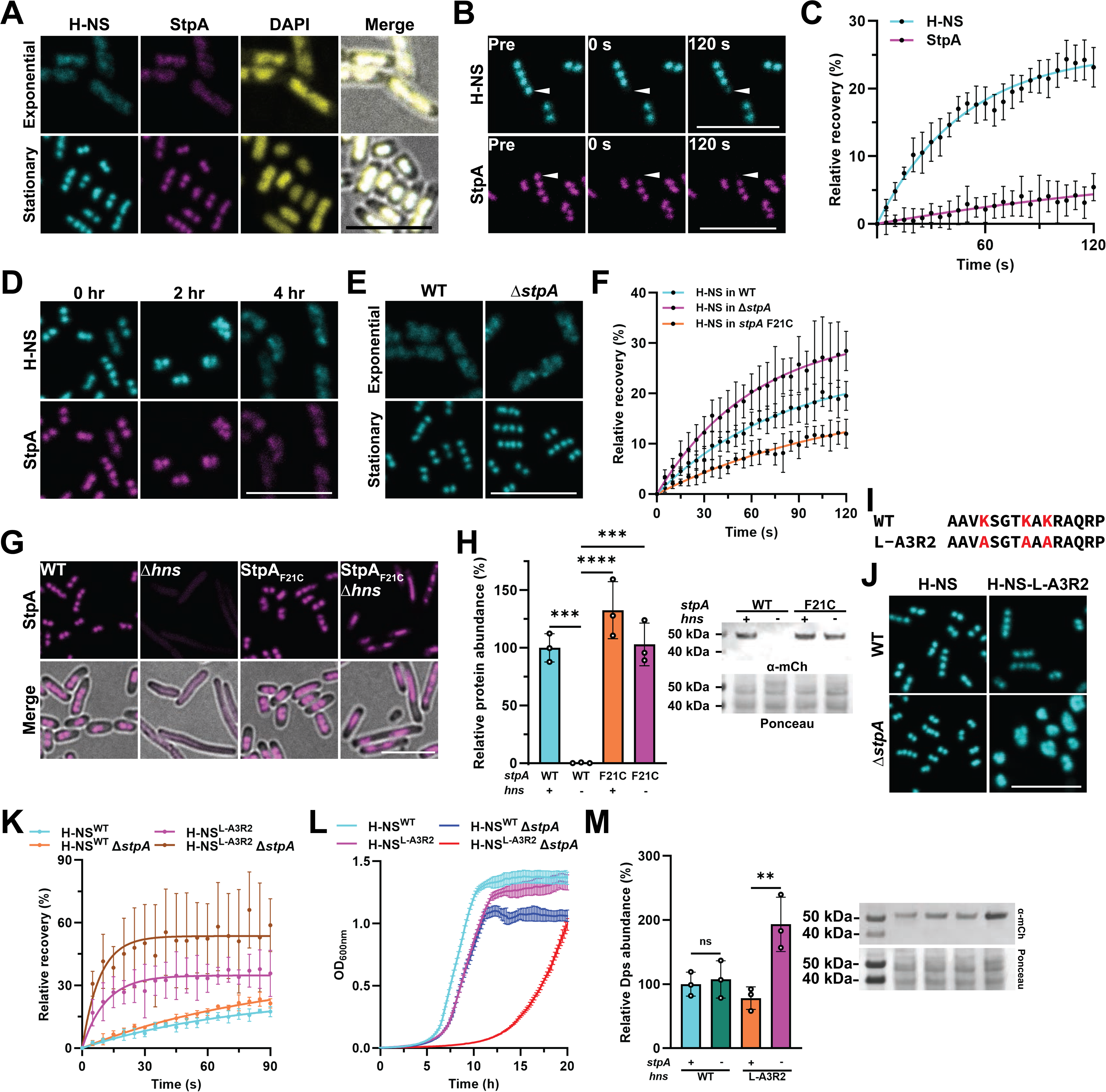
H-NS and StpA form co-condensates in *E. coli*. *E. coli* MG1655 expressing H-NS-mNG or variants (depicted in magenta) and/or StpA-mCh or variants (depicted in cyan) from their endogenous loci were grown in Gutnick+ minimal medium at 37°C until exponential or late stationary phase **A.** Live cell fluorescence images of H-NS-mNG and StpA-mCh signal obtained during exponential growth (upper panel) or late stationary phase (lower panel) are shown. Nucleoids are stained with DAPI (yellow). Overlay with brightfield (BF) is shown in the rightmost panel. **B.** Fluorescence recovery after photobleaching (FRAP) experiments on H-NS-mNG or StpA-mCh puncta contained within stationary grown cells. Frames depict puncta before (pre) or immediately upon (0 s) or 120 s after photobleaching. Arrowheads depict the photobleached area. **C.** FRAP quantification of data shown in (B) (mean, SD, *n* = 8). **D.** Fluorescence images of *E. coli* expressing H-NS-mNG and StpA-mCh before, 2 hr or 4 hr after dilution into fresh medium. **E.** Fluorescence images of H-NS-mNG expressing *E. coli* in a *stpA*^+^ or Δ*stpA* background. **F.** FRAP quantification of H-NS-mNG foci in *stpA*^+^ (cyan), Δ*stpA* (magenta) or *stpA*^F21C^ expressing *E. coli* (orange) (mean, SD, *n* = 6-8). **G.** Fluorescence images of StpA-mCh or StpA^F21C^-mCh in *hns*^+^ or Δ*hns* background. Lower panels depict the overlay of mCherry fluorescence and BF. **H.** Abundance of StpA-mCh or StpA^F21C^-mCh in *hns*^+^ or Δ*hns* strains as quantified by western blotting using antibodies against mCherry. Total protein was used as a loading control (*n* = 3, SD, one-way ANOVA, *** *p* < 0.001, **** *p* < 0.0001). A representative western blot image is shown. **I.** Linker sequence of WT H-NS or the H-NS^L-A3R2^ mutant. Mutated lysine (K) residues are highlighted in red. **J.** Fluorescence images of H-NS-mNG or H-NS^L-A3R2^-mNG in *stpA*^+^ or Δ*stpA* bacteria. **K.** FRAP analysis of H-NS-mNG signals in bacteria shown in (K) (mean, SD, *n* = 6). **L.** Growth curves of the indicated strains in Gutnick+ minimal media. Strains expressed the untagged versions of the depicted genes. **M**. Abundance of Dps-mCh in the indicated strains as quantified by western blotting against mCherry. Total protein was used as loading control (*n* = 3, SD, one-way ANOVA, ** *p* < 0.01). A representative western blot image is shown. All scale bars are 5 µm.

### H-NS and StpA reciprocally affect each other’s *in vivo* phase behaviors

To investigate whether H-NS and StpA differ in their individual phase behavior *in vivo*, we expressed H-NS-mNG in an *E. coli* Δ*stpA* strain and, conversely, StpA-mCh in a Δ*hns* strain. We did not detect any major differences in the spatial distribution of H-NS-mNG in exponentially or stationary phase Δ*stpA* strains as compared to WT *E. coli* (Fig. 1E), indicating that the formation of *in vivo* H-NS-mNG foci can occur independently of StpA. FRAP analysis of H-NS- mNG foci formed in the stationary phase Δ*stpA* strain, however, revealed an increase in their relative recovery rate compared to similar sized foci formed in the presence of StpA (Fig. 1F, compare cyan and purple traces). These results suggested that the co-presence of StpA in the condensates might reduce the dynamic diffusivity of H-NS-mNG. In contrast, in exponentially growing Δ*hns* bacteria, most StpA was found to be insoluble (Fig. S1N) and the fraction of StpA- mCh that formed discrete foci showed no significant recovery upon photobleaching (Fig. S1H, G). Moreover, in stationary phase Δ*hns* bacteria, no focal enrichment of StpA-mCh was detected (Fig. 1G). This result was not unexpected as StpA is degraded by the Lon protease in the absence of H-NS, particularly during stationary phase^38^ (Fig. 1H). To determine whether StpA has the intrinsic propensity to form condensates when it is allowed to accumulate in stationary phase, we introduced an F21C mutation (StpA^F21C^-mCh), which has previously been shown to prevent Lon-mediated degradation^38^. Expression of this variant from the endogenous *stpA* locus produces slightly higher than normal StpA levels in the presence of H-NS and WT-like levels in Δ*hns* (Fig. 1H). Yet, despite this restoration in cellular abundance, StpA^F21C^-mCh still failed to assemble into spherical puncta in the absence of H-NS (Fig. 1G). That this effect was not mutation specific became evident in H-NS-mNG expressing bacteria, where StpA^F21C^- mCh formed WT-like co-condensates (Fig. 1G). Of note, however, we did observe that the fluorescence recovery rate of H-NS-mNG in the H-NS-mNG/StpA^F21C^-mCh co-condensates was lower compared to the recovery rate seen in WT bacteria (Fig. 1F, orange trace). Combined with the increased FRAP recovery of H-NS-mNG in the Δ*stpA* strain, this finding suggested that the cellular StpA to H-NS ratio might negatively affect the dynamic fluidity of H-NS in co-condensates (compare Fig. 1F with Fig. S1O). Together, these results demonstrated that despite their high sequence homology and structural similarity, H-NS and StpA differ drastically in their *in vivo* phase behaviors and appear to reciprocally affect each other’s *in vivo* properties. Whereas H-NS phase separates on its own but is affected in its dynamic diffusivity by StpA, the paralog StpA is insoluble on its own and requires the presence of H-NS to form soluble *in vivo* condensates.

### Condensate formation appears crucial for proper gene silencing and bacterial growth

Much effort has been spent in trying to determine the physiological roles of condensate formation *in vivo*^43^. Achieving this goal, however, requires identifying mutant variants with defects in phase separation ability but minimal changes in their structure or function in the dilute phase. Given the critical role that charged residues in IDRs play in protein phase separation^44^, we now reasoned that the previously characterized linker mutant of H-NS (H-NS^L-A3R2^) might fulfill these criteria. In H-NS^L-A3R2^, three of the five positively charged IDR residues (K83, K87 and K89) are replaced with alanines (Fig. 1I). The two remaining positively charged amino acids, i.e., R90 and R93 are left intact to minimize any potential impact on DNA binding.

Importantly, this mutant variant had been previously shown to form WT-like nucleoprotein filaments^45^, indicating that it remains able to interact with itself as well as with DNA. Indeed, stationary phase *E. coli* expressing the H-NS^L-A3R2^-mNG variant from the native *hns* locus formed WT-like H-NS condensates (Fig. 1J, upper panel), albeit with significantly faster FRAP recovery rates than the WT protein (Fig. 1K, compare cyan and purple traces). Strikingly, however, and in stark contrast to H-NS-mNG, which does not require StpA for *in vivo* foci formation (Fig. 1E), this mutant H-NS variant was now dependent on the presence of StpA to form discrete foci in stationary phase bacteria (Fig. 1J, lower panel). Moreover, the FRAP recovery of regions enriched for the mutant variant in the Δ*stpA* strain was even higher than in the presence of StpA (Fig. 1K, brown traces). These results suggested that the replacement of three positively charged residues in H-NS’s linker hamper phase separation presumably by increasing the dynamic properties of the IDR. Moreover, they implied that StpA’s stabilizing effects on the fluidity of H-NS within condensates is sufficient to overcome the inability of the H- NS^L-A3R2^ variant to phase separate. These strains, which significantly differed in their capacity to form H-NS condensates, provided us now with the unique opportunity to directly test the physiological consequences of proper heterochromatin condensate formation. Analysis of their growth in minimal medium revealed that while bacteria expressing the H-NS^L-A3R2^ variant in the presence of StpA (and hence form WT-like condensates) reached almost WT-like growth (Fig. 1L, cyan *versus* purple traces), growth in the absence of StpA, which impaired condensate formation, was significantly compromised (Fig. 1L, red traces). At the same time, we observed a dramatic de-repression of the prototypical H-NS target gene *dps*^46^ upon loss of *stpA* in bacteria expressing the H-NS^L-A3R2^ variant but not in Δ*stpA* mutants expressing WT H-NS (Fig. 1M). Based on these data, we concluded that proper condensate formation plays a physiologically relevant role in H-NS/StpA mediated gene silencing and *E. coli* growth.

### H-NS and StpA display highly distinct phase behaviors *in vitro*

To investigate why H-NS and StpA behave so differently *in vivo*, we purified both proteins and investigated their *in vitro* phase behaviors under physiologically relevant buffer conditions^47^. For visualization and FRAP experiments, we supplemented H-NS with 1-4% of Cy3-labeled H-NS and StpA with 1-4% of Cy5-labeled StpA^F21C^. Supplementation with the labeled proteins did not detectably affect the phase behavior and both labeled and unlabeled protein preparations were used interchangeably. We found that upon dilution from their high salt storage buffers into the phase separation buffer, H-NS readily assembles into spherical condensate-like droplets (Fig. 2A, upper panel) whereas StpA, at concentrations as low as 1.25 µM, precipitates in form of seemingly amorphous aggregates (Fig. 2A, lower panel). Co-incubation of H-NS with a FAM-labeled 150 bp DNA sequence amplified from an endogenous H-NS binding site^14^ caused the formation of heterotypic H-NS/DNA co-condensates (Fig. 2B, upper panel) and a reduction in the H-NS saturation concentration (*c*_sat_) from 25 µM to 5 µM (Figs. 2C, S2A). Unexpectedly, we found that FAM-DNA also colocalized with the amorphous aggregates of StpA (Fig. 2B, lower panel), suggesting that precipitated StpA retains its DNA binding activity. Subsequent FRAP analysis of homotypic H-NS condensates revealed that the Cy3-H-NS fluorescence rapidly recovered upon both partial- (Fig. 2D) and full-droplet (Fig. S2B) photobleaching and seemed unaffected by the presence of DNA (Figs. 2E, S2C). No significant recovery was detected in the case of StpA, independent of the absence or presence of DNA (Figs. 2D, F). To explore the structural properties of these assemblies in more detail, we performed cryo-electron tomography (cryo-ET) analysis on H-NS condensates and transmission electron microscopy (TEM) on the insoluble StpA precipitates. Cryo-ET analysis revealed a dense mixture of linear structures in H-NS condensates (Fig. 2G), consistent with previous studies showing that H-NS dimers form short linear oligomers^48,49^. In the presence of DNA, the condensates displayed no obvious differences in their coarse morphologies (Fig. 2G). Unexpectedly, however, TEM analysis of the insoluble StpA assemblies also revealed linear structures, yet apparently thicker and longer than those observed in H-NS condensates (Figs. 2H, I). As was the case for H-NS, presence of DNA did not cause any significant differences in the fibrillar structure of StpA. Of note, no such fibrils were observed for H-NS under the same conditions (Fig. S2D). Taken together, these data provided good *in vitro* support for our *in vivo* findings, which showed that on their own, the two closely related paralogs H-NS and StpA differ drastically in their solubility and phase behavior.

**Figure 2.**
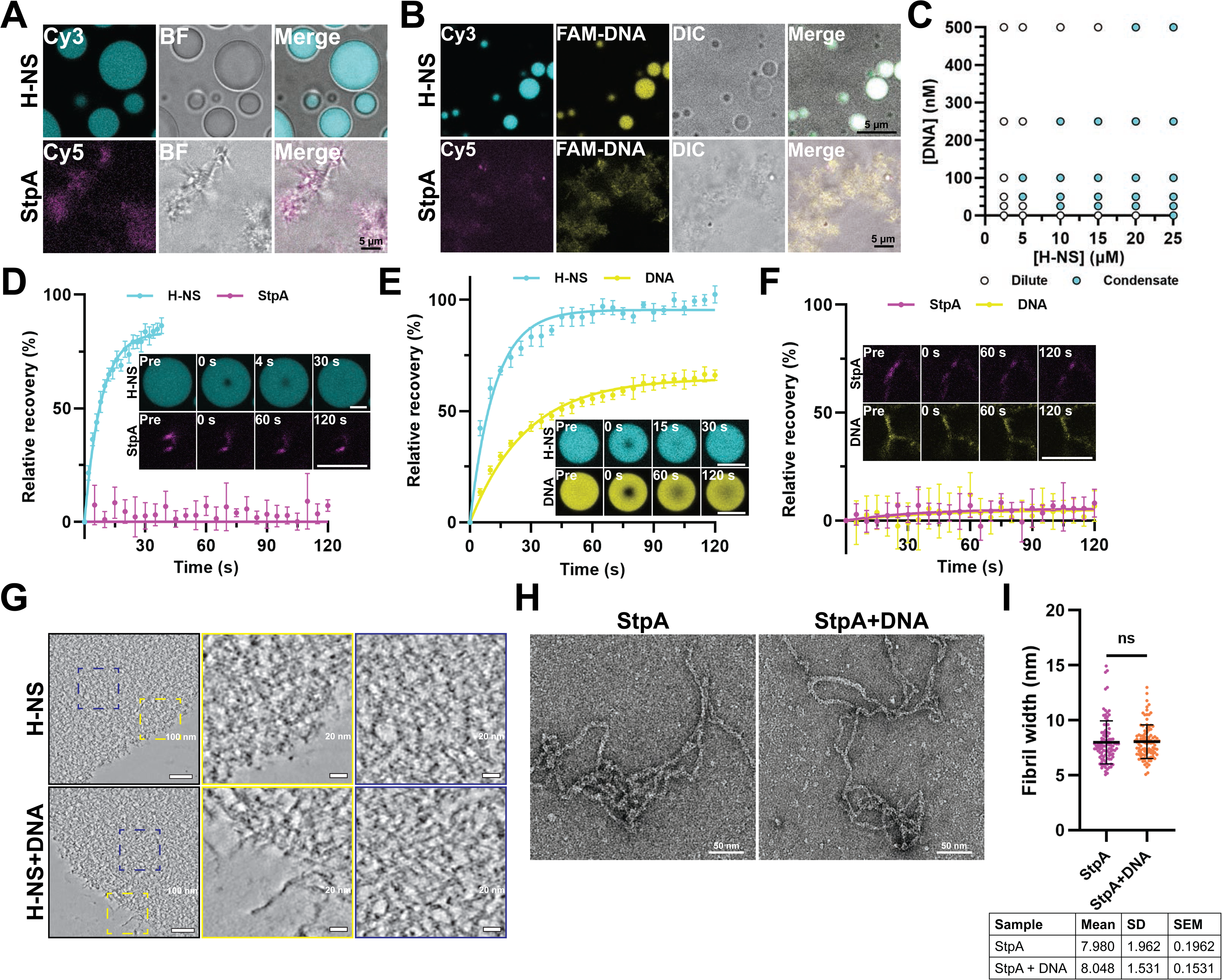
H-NS and StpA differ in their *in vitro* phase behavior. **A.** Fluorescence and bright field (BF) images of 100 µM purified H-NS (+ 1% Cy3-H-NS) or 100 µM purified StpA (+ 1% Cy3-StpA^F21C^) upon dilution from high salt buffer into GMT-buffer (100 mM Na-Glu, 1 mM MgAc, 20 mM Tris-Ac, pH 7.5). Residual NaCl concentration was 30 mM (H-NS) or 50 mM (StpA), respectively. Images were taken after 20 minutes incubation at room temperature unless otherwise indicated. **B.** Same as in (A) except that 25 µM H-NS (+ 4% Cy3-H-NS) or 25 µM StpA (+ 4% Cy5-StpAF21C) in the presence of 100 nM FAM-labeled 150 bp DNA was used. **C.** Phase diagram of H-NS and 150 bp DNA in GMT buffer based on microscopy (see Fig. S2A). **D.** Quantification of partial FRAP analysis of H-NS (cyan) droplets or StpA (magenta) precipitates shown in (A); mean, SD, *n* = 4-6. **Inset:** Representative images of FRAP time course. **E.** and **F.** Quantification of partial FRAP of H-NS/DNA droplets (E) or StpA/DNA (F) precipitates shown in (B); color code as before; DNA shown in orange (mean, SD, *n* = 4). **Inset:** Representative images of FRAP time course. **G.** Cryo-ET images of condensates composed of 50 µM H-NS or 25 µM H-NS with 100 nM DNA. Inset in the left panel are zoomed in on the right. **H.** TEM image of insoluble 2.5 µM StpA precipitates in absence or presence of 5 nM DNA. **I.** Analysis of the mean width of StpA fibrils shown in (H) (*t* test, *n* = 100). Scale bars are 5 µm unless otherwise indicated.

### H-NS converts insoluble StpA fibrils into liquid condensates

Our *in vivo* results revealed a strong dependence of StpA solubility on the presence of H-NS. To investigate whether H-NS affects also the *in vitro* solubility of StpA, we incubated pre-formed insoluble StpA fibrils with either H-NS or DNA and H-NS. To our surprise, in either case we observed the immediate formation of mixed precipitates, followed by their slow conversion into spherical co-condensates (Figs. 3A-B; S3A-B; movie 1). Once the droplets formed, both H-NS and StpA displayed significant dynamic diffusivity within these droplets (Figs. 3C, S3C), suggesting that these compartments are indeed heterotypic liquid condensates. Moreover, and as shown before *in vivo*, the relative fluidity of StpA remained significantly lower compared to H-NS while the fluidity of DNA appeared to follow that of StpA (Figs. 3C, S3C). A phase diagram of H-NS *versus* StpA generated after 3 hours of incubation demonstrated that i) H-NS concentrations as low as 1.25 µM convert 20 µM of insoluble StpA assemblies into liquid condensates and ii) that co-incubation of 1.25 µM pre-formed StpA-fibrils and 1.25 µM H-NS is sufficient to form mixed droplets, effectively reducing the *c*_sat_ of H-NS by a factor of 20 (Fig. 3D). These results suggested that StpA, once in complex with H-NS, contributes features that promote their phase separation. Dissolution of StpA fibrils was dependent on the presence of H-NS and independent of DNA (Fig. 3E). Yet, the rate at which these co-precipitates converted into co-condensates was sensitive to the H-NS concentration (Fig. S3D) and fastest when H-NS/DNA complexes were added to pre-formed StpA fibrils (Fig. 3E). Of note, when we added H-NS to pre-formed StpA fibrils at concentrations far above its own *c*_sat_, H-NS immediately formed large condensates that completely engulfed the StpA assemblies (Fig. 3F). Within minutes, the StpA assemblies dissolved inside these condensates, giving rise to homogeneous binary condensates. Of note, Hfq, an unrelated NAP with similar *in vitro* phase separating propensities as H-NS^50^ was entirely ineffective in dissolving StpA precipitates (Fig. S3E). These results suggested that the ability to trigger StpA phase transition is H-NS specific. Moreover, analysis of the H-NS^L-A3R2^ mutant, which, as described above, forms *in vivo* foci in the presence of StpA but not in its absence, lacked the capacity to phase separate on its own (Fig. S3F). Yet, the variant dissolved pre-formed StpA fibrils (Fig. S3G) and formed heterotypic condensates with StpA that displayed dynamic diffusivity like WT H-NS/StpA/DNA condensates (Figs. S3H, I).

**Figure 3.**
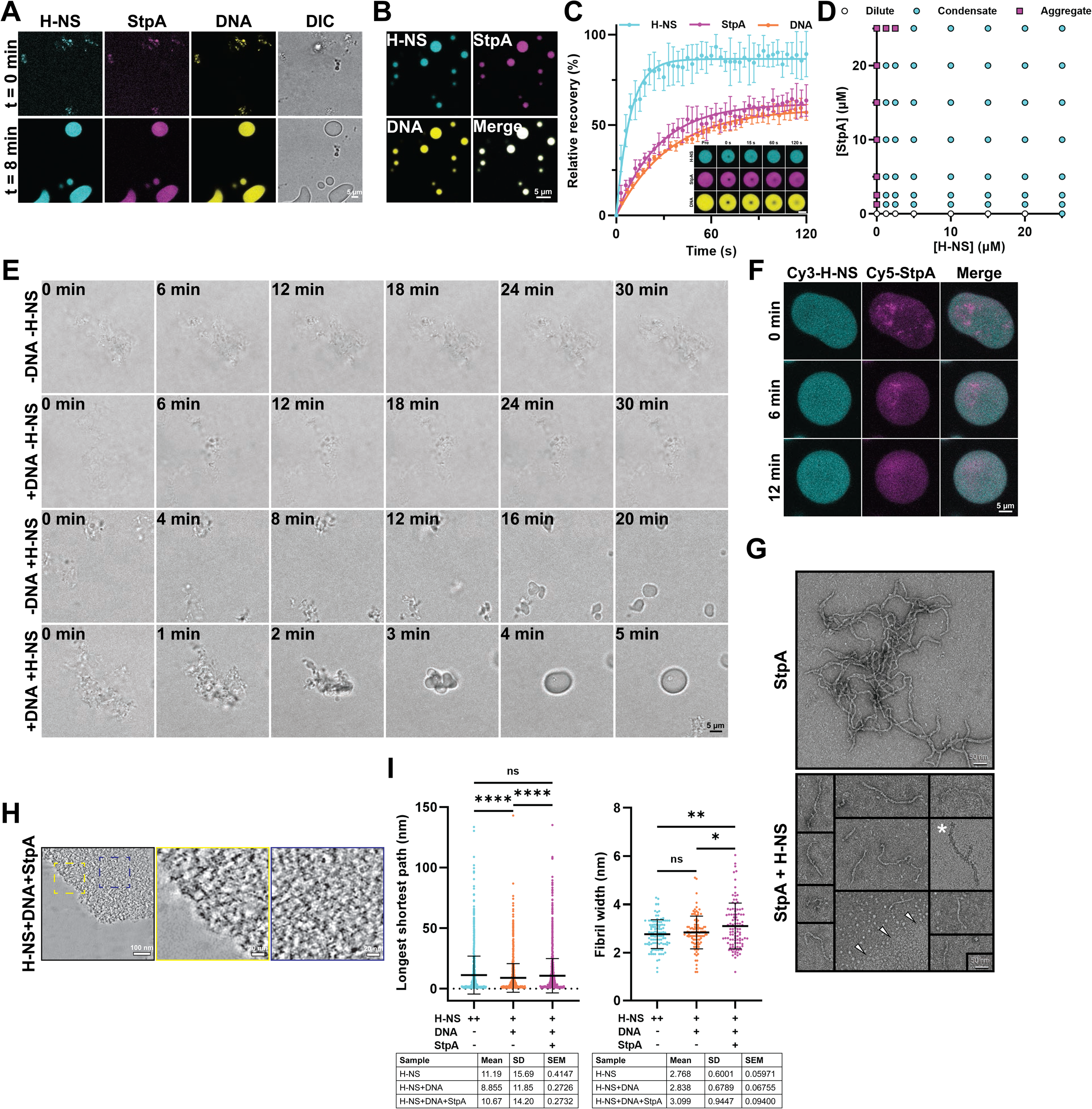
H-NS and StpA affect each other’s phase behavior *in vitro*. **A.** Fluorescence and BF images of 25 µM H-NS (+ 4% Cy3-H-NS), 10 µM preformed StpA (+ 4% Cy5-StpA^F21C^) fibrils and 100 nM FAM-DNA immediately upon and 8 min after mixture in GMT buffer. **B.** Fluorescence images of the same samples as in (A) after 30 min of incubation. **C.** Quantification of partial FRAP of H-NS/StpA/DNA co-condensates shown in (B). Color code as before (mean, SD, *n* = 4-6). **Inset:** Representative images of FRAP time course. **D.** Phase diagram of H-NS and StpA acquired 3h after incubation in GMT buffer. NaCl concentration across all samples was adjusted to 40 mM. **E.** BF images of 10 µM StpA in GMT buffer at indicated time points after addition of 25 µM H-NS and/or 100 nM DNA. **F.** Fluorescence images of 50 µM H-NS (+ 2% Cy3-H-NS) and 10 µM preformed StpA (+ 4% Cy5-StpA^F21C^) fibrils immediately, 6 min or 12 min after mixing in GMT buffer. **G.** TEM images of 2.5 µM StpA without or with 2.5 µM H-NS after 30 min of incubation in GMT buffer. Arrowheads indicate boundary of presumed condensates. Asterisk indicates fibril embedded within presumed condensates. **H.** Cryo-ET images of condensates composed of 25 µM H-NS, 10 µM StpA and 100 nM DNA. Inset in the left panel are zoomed in on the right. **I.** Quantification of linear structure length (longest shortest path) and fibril width inside condensates (length *n* = 143-270, width *n* = 101, one-way ANOVA, * *p* < 0.05, ** *p* < 0.01, **** *p* < 0.0001). Scale bars: 5 µm, unless otherwise indicated.

The observed conversion of mesoscale StpA precipitates into liquid condensates prompted us to investigate how, at the molecular level, H-NS might modify StpA. TEM analysis of pre-formed StpA fibrils revealed that the addition of H-NS dramatically decreased their length, eventually causing their disappearance (Fig. 3G). Short StpA fibrils were even observed within protein-dense areas that presumably correspond to condensates (Fig. 3G, asterisk). Unexpectedly, however, cryo-ET analysis of the mixed condensates did not reveal any gross morphological differences between the linear oligomers observed in H-NS, H-NS/DNA or H-NS/StpA/DNA condensates (Figs. 2G and 3H). Due to the high density of oligomers in condensates, we were unable to conduct any detailed structural analysis. We therefore decided to compare the width and length of linear oligomers in all three types of condensates by skeletonizing the linear structures taken from the central slice of each tomogram (Fig. 3I)^51^, as well as to compare oligomer width. These analyses revealed that the oligomers in the H-NS/StpA/DNA condensates are slightly but significantly longer and wider than the oligomers of H-NS/DNA or H-NS condensates (Fig. 3I). Together, these findings recapitulated our *in vivo* observations and demonstrated that H-NS can convert insoluble StpA fibrils into shorter fibrils that eventually co-phase separate with H-NS. They also revealed StpA’s reciprocal impact on H-NS, in which StpA, potentially through alterations in the oligomer structures, modulates the phase landscape of its paralog.

### StpA to H-NS ratio defines fluidity, thermal stability and salt resistance of droplets

Based on these results, we now wondered whether interacting paralogs such as H-NS and StpA, through their distinct phase behaviors, might provide organisms with the capacity to adjust the dynamic properties and stability of condensates over a wider range of environmental conditions. We found this idea intriguing as previous studies demonstrated that StpA concentrations go up relative to H-NS levels with increasing temperature or osmolarity^24,40^. To directly test this hypothesis, we generated mixed H-NS/StpA/DNA droplets with constant total protein (25 µM) and DNA (100 nM) concentrations and variable StpA to H-NS ratios. Droplets were clearly noticeable in all samples except the StpA alone sample. Subsequent FRAP analysis revealed that as we increased the ratio of StpA to H-NS from 1:10 to 10:1, the rates and yields of fluorescence recovery significantly decreased for all three condensate components (Figs. 4A, B). These results supported the idea that the relative ratio between StpA and H-NS defines the diffusivity of both proteins and DNA in the condensates in agreement with our earlier *in vivo* observations (Fig. 1F). Dynamic diffusivity and stress resistance of condensates are both regulated by the type and strength of multivalent interactions in the system and often display a negative correlation^52–54^. Therefore, we next tested the stability of our condensates by either increasing the osmolarity of the solution or the environmental temperature. Whereas incubation of H-NS condensates in 100 mM NaCl fully dissolved the droplets, StpA addition at 1/10 of the H-NS concentration was sufficient to maintain droplet formation (Fig. S4A). NaCl titration experiments confirmed that by increasing the StpA:H-NS ratio, the mixed condensates, both with (Figs. 4C) and without DNA (Fig. S4B), became increasingly more salt resistant, eventually reaching the NaCl resistance of pure StpA fibrils (i.e., 500 mM NaCl). The thermal stability of the droplets was equally dependent on the presence of StpA in the mixture (Fig. 4D). Whereas H-NS/DNA droplets dissolved upon incubation at 45°C, presence of StpA appeared to maintain droplet stability (Fig. 4D). StpA/DNA mixtures still failed to phase separate on their own even at these elevated temperatures (Fig. 4D). These results provided *in vitro* support for a model by which organisms evolved paralogs such as the H-NS/StpA pair, which are located at the apparently opposite ends of the phase spectrum, to maintain condensate formation, fluidity, stability and likely function under a wide range of environmental stress conditions.

**Figure 4.**
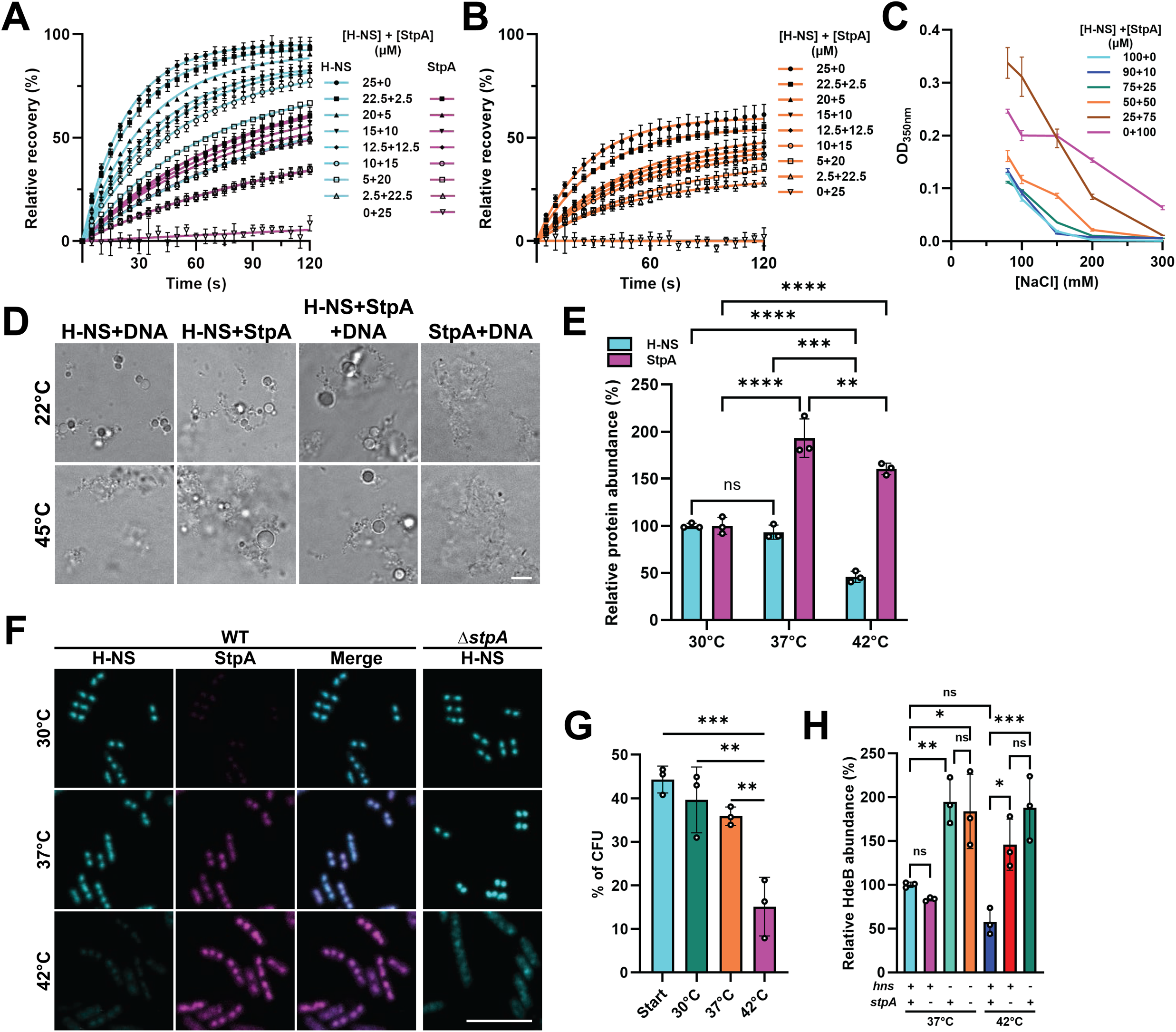
StpA to H-NS ratio defines condensate properties and bacterial fitness. **A.** Quantification of full FRAP of H-NS and StpA in H-NS/StpA/DNA co-condensates. The total protein and DNA concentrations were constant at 25 µM and 100 nM, respectively. The NaCl concentration across all samples was adjusted to 80 mM. **B.** Quantification of partial FRAP of DNA in the same H-NS/StpA/DNA co-condensates shown in (A). **C.** Turbidity measurements of H-NS/StpA/DNA mixtures measured at different NaCl concentrations. Concentrations of H-NS and StpA are indicated. DNA concentration was kept constant at 100 nM. **D.** BF images of 25 µM H-NS and/or 10 µM StpA with or without 100 nM DNA in GMT buffer. Samples were either incubated at 22°C for 40 minutes or at 22°C for 20 minutes and then at 45°C for another 20 minutes. Before imaging, samples were fixed with 4% formaldehyde to preserve condensate morphology. **E.** Abundance of endogenously expressed H-NS-mNG and StpA-mCh in *E. coli* grown for 24h at the indicated temperatures. Quantification was performed by western blotting against mNG (H-NS) or mCh (StpA). Ponceau S staining of total protein was used for normalization; one-way ANOVA, *n* = 3, ** *p* < 0.01, *** *p* < 0.001, **** *p* < 0.0001). **F.** Fluorescence images of *E. coli* endogenously co-expressing H-NS-mNG and StpA-mCh, or H-NS-mNG only in a Δ*stpA* background grown for 24h at indicated temperatures. **G.** Percentage of *stpA::Kan* strain CFU over total CFU before and 24h after co-culture with WT *E. coli* at the indicated temperatures. Mean, SD, one-way ANOVA, *n* = 3, ** *p* < 0.01, *** *p* < 0.001. **H.** Abundance of endogenously expressed HdeB-3xFLAG in *E. coli* MG1655 WT, Δ*hns*, Δ*stpA* and Δ*hns*Δ*stpA* strains, grown to mid-exponential phase at 37°C or 42°C. No growth for the Δ*hns*Δ*stpA* strain was observed at 42°C (Fig. S4C). Quantification was performed by western blotting using antibodies against the 3xFLAG tag. Ponceau S staining of total protein was used for normalization (one-way ANOVA, *n* = 3, * *p* < 0.05, ** *p* < 0.01, *** *p* < 0.001). Scale bars: 5 µm.

### Stress-induced StpA:H-NS changes maintain *in vivo* condensate stability and function

To directly test this model *in vivo*, we next monitored temperature-induced changes in relative StpA and H-NS levels using H-NS-mNG and StpA-mCh expressing *E. coli* grown between 30°C to 42°C. Western blot analysis using antibodies against the fluorescent tags revealed that the relative StpA:H-NS ratio increased about 4-fold as we increased the growth temperature from 30°C to 42°C (Fig. 4E). This change in ratio was also reflected in the H-NS/StpA foci formed in stationary phase, which contained increasingly more StpA relative to H-NS as the growth temperatures was raised (Fig. 4F). Our observation that at 42°C, H-NS-mNG no longer formed discrete foci when *stpA* was deleted while H-NS foci formation at non-stress temperatures was independent of StpA (Fig. 1E) provided first *in vivo* evidence in support of our model (Fig. 4F).

Subsequent analysis of growth rates further confirmed this temperature-dependence on StpA by revealing that an increase in growth temperature causes an increasingly more severe growth defect in the Δ*stpA* strain (Fig. S4C). Moreover, when we co-cultured WT bacteria with equal colony forming units (CFUs) of the *stpA::Kan* strain for 24h, we observed a drastically reduced CFU% of the *stpA* deletion strain at 42°C but not lower temperatures (Figs. 4G and S4D), indicating that StpA becomes increasingly more important for maintaining competitive fitness at high temperatures. To link these defects with the endogenous functions of H-NS/StpA, we tested whether StpA expands the temperature range of transcriptional silencing. We tested bacterial strains endogenously expressing known gene targets of H-NS (*i.e.*, *dps*, *gadA*, *hdeB*)^46^, epitope tagged to facilitate their detection via immunoblotting. This approach is superior to qPCR as it eases normalization and is independent of the identification of appropriate housekeeping genes that are unaffected by growth conditions. We found that the expression of all three tested targets was de-repressed upon loss of *stpA* at 42°C but not 37°C (Figs. 4H, S4E, F), suggesting that under high temperatures, H-NS requires StpA to support gene silencing. We made very similar observations when we exposed our bacterial strains to osmotic stress. As we increased the NaCl levels in our growth medium, H-NS-mNG expressing bacteria became increasingly dependent on the presence of StpA for foci formation (Fig. S4G), and more reliant on StpA to maintain their growth rates (Figs. S4G, H) and competitive fitness (Fig. S4I). Together, these data provide evidence that StpA provides bacteria with improved resistance and fitness across a broad range of adverse growth conditions likely through its ability to stabilize H-NS in mixed nucleoid-associated condensates.

### Structural differences in H-NS/StpA homo-and hetero-oligomers

The distinct behaviors of H-NS and StpA prompted us to search for potential structural differences between the paralogs. Structural modeling of *E. coli* H-NS/StpA dimers and tetramers built using Chai discovery or using AlphaFold 3^55^ confirmed the presence of two well-defined interfaces - an antiparallel coiled-coil dimerization site 1 (D1) and a helix-turn-helix dimerization site 2 (D2) in the N- and C-terminal portions of the oligomerization domain, respectively (Fig. S5A)^48^. D1 mediates the formation of intertwined “head-to-head” dimers, whereas interactions between D2 promote tail-to-tail polymerization and give rise to a linear chain of H-NS molecules^48,56^. Previous studies suggested that while the D1 in H-NS is thermostable and resistant towards high osmolarity, D2 is temperature and salt sensitive^56–58^, a result that agreed well with our observation that H-NS condensates are temperature and salt labile. To understand the observed stability differences between H-NS and StpA homo-oligomers, we performed all-atom molecular dynamics (MD) simulations on both proteins. As a proof of concept, we first conducted MD simulations using the truncated H-NS (aa 1-82) and StpA (aa 1-83) homo-dimers. These experiments confirmed that the interactions at D1 are stable under hyperosmotic conditions (500 mM KCl at 37°C) in both proteins (Fig. S5B).

Subsequently, we compared the dynamics of full-length H-NS/StpA homo- and hetero-tetramers under the same hyperosmotic conditions using the AlphaFold-model structures as input (Fig. S5C). Overall backbone Cα root-mean-square deviation (RMSD) of both H-NS and StpA homo- tetramers drifted towards the end of simulation, indicating high conformational plasticity (Fig. S5D). In contrast, H-NS/StpA hetero-tetramers adopted a low plateau RMSD, suggesting a more rigid compromise geometry. A more detailed structural analysis highlighted these differences. While the H-NS homo-tetramer retained a relatively linear organization (Fig. 5A), the StpA homo-tetramer adopted a pronounced curvature (Fig. 5B). The hetero-tetramer, in comparison, appeared more linear than StpA alone, consistent with its lower RMSD values and suggestive of dampened flexibility (Fig. 5C). The significant curvature in StpA homo-tetramers was rooted in D2, where the helices engaged in symmetric interactions via three charged side chains (Lys57, Arg62, Asp68). This interface was significantly more flexible in H-NS homo- tetramers and uniformly more stable in StpA and StpA/H-NS tetramers as indicated by their lower root-mean-square fluctuation (RMSF) across the same residues (Figs. 5D-F). Most unexpectedly, however, we observed that in both StpA tetramer and mixed tetramers, the DNA binding domain (DBD) of one dimer engaged in stable and symmetric interactions with the coiled-coil surface of D1 in the neighboring dimer (Fig. 5B and 5C). This contrasted with H-NS, where the DBD flexibly radiates outward, only forming very transient interactions with its own D1 (Fig. 5A)^56^. Together, these data suggested that while both paralogs formed stable dimers through D1, StpA engages in more stable homo- and heterotypic oligomers through the symmetric interactions not only at D2 but also between the DBD and D1, effectively interlocking the adjacent dimers. To experimentally validate these results and determine the contributions of individual structural domains to their phase behavior, we domain-swapped the respective oligomerization domains, generating the chimeras H-S-S and S-H-H (Fig. 5G). The S-H-H chimera was extremely insoluble and could not be purified, implying that the oligomerization domain is primarily responsible for StpA’s insolubility. The H-S-S chimera, in contrast, behaved like an improved versions of WT H-NS, forming biomolecular condensates at a significantly lower *c*_sat_ (5 µM), which were only mildly affected by the additional presence of StpA (Fig. 5H) and seemingly unaffected by DNA (Fig. S5E). These results suggested that the oligomerization domain was the main determinant of the proteins’ phase behaviors while StpA’s linker and DBD, presumably through their unique ability to interact with D1, work in *trans* to enhance phase separation. In summary, these studies suggest that the N-terminal oligomerization domain defines the phase behavior of the paralogs, by causing H-NS to form condensation-competent short oligomers and StpA to form insoluble long fibrils. StpA facilitates stable homo- and heterotypic tail-to-tail interactions via D2, allowing StpA fibrils and H-NS/StpA co-condensates to remain unperturbed upon stress. Finally, inter-dimer interactions between DBD and D1 promote additional support to oligomer stability and help to fine-tune the formation and dynamics of co- condensates.

**Figure 5.**
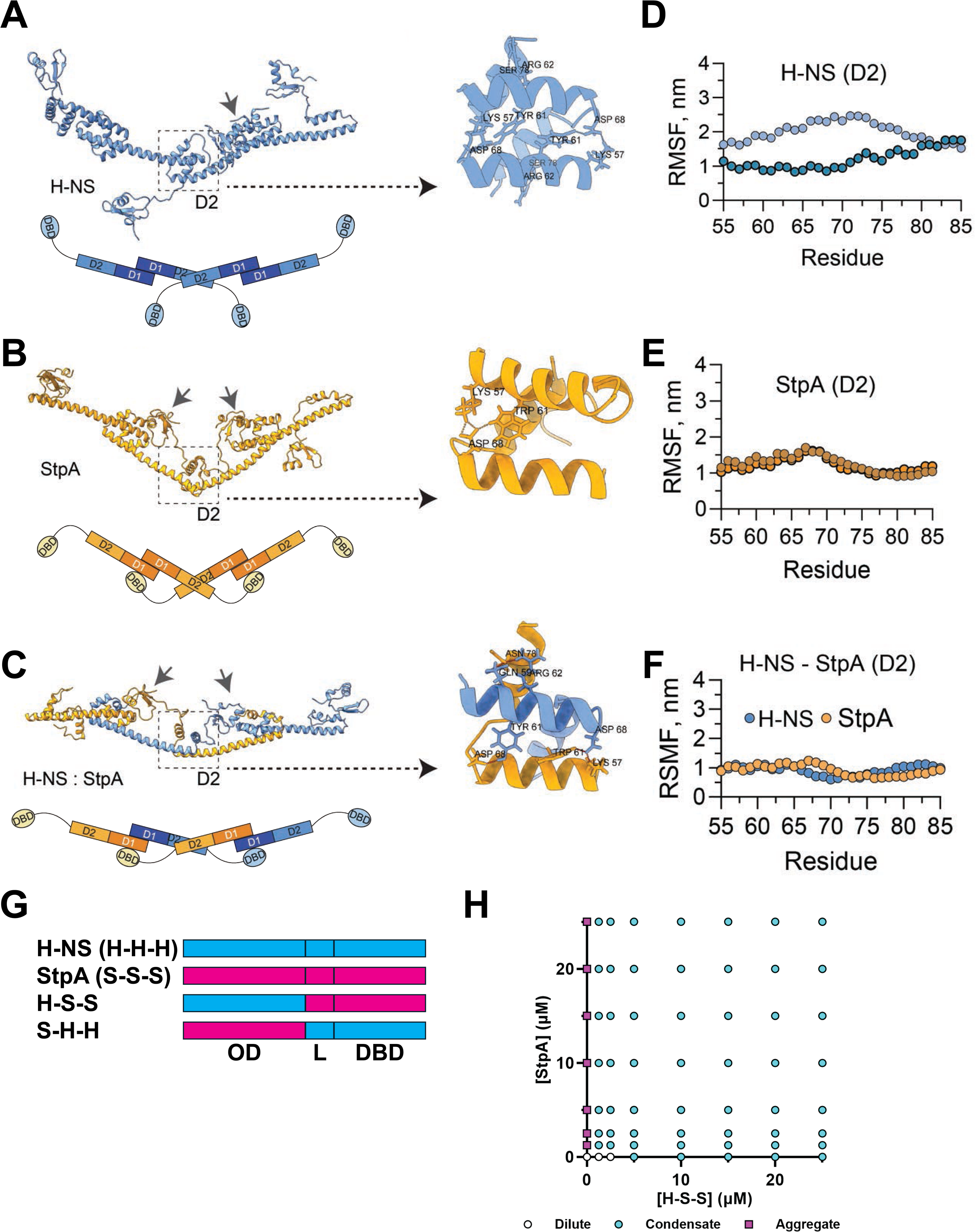
H-NS and StpA oligomers display distinct structural features. **A-C.** Final 200 ns snapshots of (A) H-NS tetramer, (B) StpA tetramer or (C) 2:2 H-NS:StpA tetramers modeled in 500 mM KCl at 37°C. Arrows highlight the DBD forming transient interactions with dimerization site 1 (D1). Zoom-in images highlight dimerization site 2 (D2) interface with residues involved in oligomerization. A model of H-NS oligomers is depicted below each simulation. H-NS and StpA molecules are colored as blue and gold, respectively. **D-F.** Root-mean-square fluctuations (RMSF) of residues spanning the D2 interface in (D) H-NS tetramer, (E) StpA tetramer or (F) 2:2 H-NS:StpA tetramer. **G.** Model demonstrating domain organization in H-NS, StpA and the domain-swapping chimeras. **H.** Phase diagram of H-S-S chimera and StpA after 3h incubation in GMT buffer. NaCl concentration across all samples were adjusted to 40 mM.

## DISCUSSION

### A novel stress paradigm for condensate dynamics, stability and function

Over recent years, countless condensate-forming scaffolding and client proteins have been identified, and the number of biological processes that involve the formation of phase-separated condensates is continuously expanding^59–61^. However, it is still unclear how organisms adjust the stability of these condensates in response to changing environmental conditions, such as during osmotic or heat stress. These stress conditions are known to interfere with the multivalent interactions that stabilize condensates and have been shown to readily dissolve protein condensates *in vitro*^52,62–64^, making it important to solve this problem. Previous studies suggested the involvement of chaperones^65^ or posttranslational modifications^66,67^ in preventing condensate dissolution or aggregation. In this study, we provide evidence for a fresh perspective. We demonstrate that organisms facilitate condensate formation under a variety of environmental stress conditions by stoichiometrically adjusting the concentration of protein paralogs with distinct phase behaviors. We show that in the case of the NAP paralogs H-NS and StpA, this adjustment enables *E. coli* to dynamically balance the fluidity and environmental stability of condensates that constitute functional heterochromatin, thereby establishing effective gene silencing across a wide range of environmental conditions (Fig. 6A). Under physiological conditions, H-NS forms short linear oligomers that undergo phase separation, giving rise to biomolecular condensates associated with bacterial functional heterochromatin (Fig. 6B). These condensates are metastable and dissolve upon heat, osmolarity and presumably other stresses, impairing gene silencing and leading to reduced bacterial fitness^68^. In contrast, we found that the H-NS paralog StpA forms longer fibrils with higher stability and poorer solubility. Unlike ultra-stable pathological amyloid fibrils^69–71^, however, StpA fibrils can be readily degraded by bacterial proteases and converted into liquid condensates by sub-stoichiometric amounts of H-NS. Interaction between H-NS condensates and StpA fibrils yield in heterotypic condensates featuring both H-NS’s fluidity and StpA’s stability. Thus, by regulating the relative abundance of H-NS and StpA, these two features can be dynamically tuned. We show that under optimal growth conditions, StpA is kept at low levels to maintain nucleoid fluidity and gene accessibility. Upon stress, however, bacteria accumulate StpA to stabilize heterochromatin condensates, preventing uncontrolled gene de-repression.

**Figure 6.**
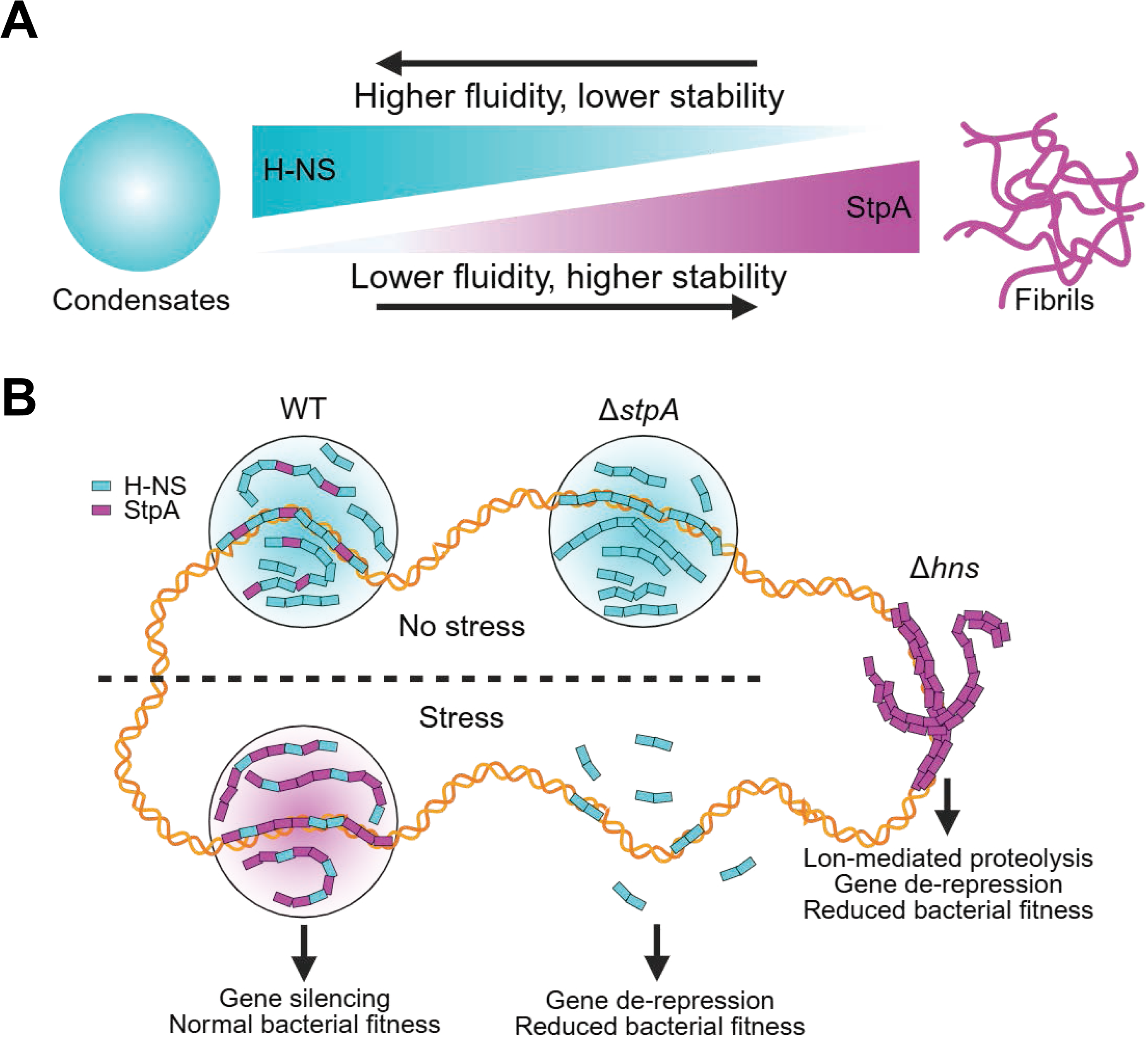
Model of bacterial heterochromatin condensate modulation by interplay between NAP paralogs. **A.** NAP paralogs H-NS and StpA lie at different ends of the phase spectrum. H-NS forms biomolecular condensates while StpA forms insoluble fibrils. **B.** Innate control of H-NS to StpA ratios under different growth conditions modulates the dynamic properties, stability and function of bacterial heterochromatin condensates.

### Differences in H-NS and StpA structure contribute to distinct phase behaviors

Our MD-simulations revealed significant structural differences between H-NS, StpA and H-NS/StpA oligomers, which likely explain their different phase behaviors. As previously reported, D2 in H-NS homo-oligomers is unstable, rendering oligomers vulnerable to stress-induced dissociation^57,58^. In contrast, the same dimerization site is involved in less dynamic interactions in StpA homo-oligomers and mixed hetero-oligomers, likely explaining their ability to remain intact during stress. Additionally, we noticed that in StpA homo- and mixed hetero-oligomers, the C-terminal DBD of one dimer forms frequent contacts with the D1 of its neighboring dimer. We speculate that such interactions, which are distinct from the previously proposed autoinhibitory conformation found in H-NS dimers^57^, facilitate the interlocking of neighboring dimers, thus further enhancing oligomer stability. We assume that similar interactions can also take place between the DBD and D1 of different oligomers, thereby contributing to the multivalency that favors biomolecular condensation^72,73^. Although our data do not provide direct evidence explaining why StpA forms insoluble fibrils while H-NS phase separates, our MD simulations reveal distinct differences in the geometry of the respective fibrils. Although more evidence is needed, such curvature could potentially facilitate the intertwining and collapse of individual oligomers into a thick filamentous structure, leading to the insoluble StpA fibrils observed in our study.

### Paralogs in biology – More support than backup systems?

The low solubility of StpA assemblies, combined with their tendency to dynamically arrest DNA, renders StpA a poor substitute for H-NS and undermines the longstanding “molecular backup” perception^36^. Instead, our results suggest that StpA serves as a support system, regulating H-NS’s phase behavior in a concentration-dependent manner. This model is also in better accordance with previous reports that StpA mutants with weaker oligomerization tendency (and presumably higher solubility) are better at compensating for the loss of H-NS^37^. It also explains the necessity of proteolytic removal of StpA in Δ*hns* bacteria, as insoluble assemblies that dynamically arrest DNA could potentially impair transcriptional response to cellular and external cues. StpA’s ability to support H-NS function makes it now reasonable to assume that such supportive roles exist for other H-NS/H-NS-like protein paralogs as well. For instance, the *H*-NS-*f*amily *p*rotein Hfp, encoded in uropathogenic *E. coli*, is transcriptionally induced by low temperature and its deletion causes a significant growth defect at 25°C^74^. Based on our study, it is tempting to speculate that other H-NS paralogs exist that support H-NS condensate formation and function in response to other environmental stress(es). By being stoichiometrically tunable, these paralogs allow precise, niche-dependent control over gene expression, and could have profound implications in organismal adaptation and survival. A similar system might also exist in eukaryotes, where biomolecular condensates are abundant and equally in need of mechanisms to withstand stress conditions that directly interfere with the forces that keep condensates intact. Earlier studies demonstrated that condensation of *h*eterochromatin *p*rotein *1* (HP1) establishes transcriptionally silent chromatin domains in eukaryotes including human, *Drosophila* and yeast^67,75–77^. These liquid compartments are suggested to organize DNA into a compact yet dynamically accessible state^78,79^. They are also responsible for the selective permeability of heterochromatin by excluding certain biomolecules from the dense phase^75^. Various HP1 paralogs feature different enrichment patterns of acidic residues in their IDRs, leading to distinct phase behaviors^80^. While HP1α and HP1γ readily form condensates with DNA, HP1β remains in a dilute state and can effectively dissolve condensates formed by its paralogs^78,81,82^. It has been proposed that while HP1α plays a central role in condensation that is supported by HP1γ, HP1β antagonizes HP1 condensation, hence limiting spreading of heterochromatin^80^. By controlling the relative abundance of HP1 paralogs, heterochromatin can thus be formed, stabilized or dissolved in a tunable manner^80^.

### Biomolecular condensate formation: A widespread mechanism for gene silencing

Prokaryotic genomes contain protein-dense and transcriptionally silent chromatin domains that are structurally distinct from but functionally analogous to eukaryotic heterochromatin^83,84^. As major constituents of bacterial functional heterochromatin^14,84^, the interplay of H-NS and StpA suggests the universal involvement of biomolecular condensates in genome organization throughout evolution. *E. coli*’s high genetic tractability also allows us to demonstrate the significance of biomolecular condensation in gene silencing and other aspects of physiology. Using an H-NS linker mutant, which is still capable of forming nucleoprotein filaments but unable to undergo condensation^45^, we demonstrated that the inability to form heterochromatin condensates leads to gene de-repression and significant growth defects. We observed similar effects when H-NS condensates could no longer form at elevated temperature or osmotic stress in the absence of StpA. These data suggest that H-NS/StpA, and likely other repressors, form nucleoprotein filaments^13,24,85^ that lead to biomolecular condensates, which, in turn, facilitate gene silencing. It is likely that H-NS and StpA molecules that are present within the condensates but not directly bound to DNA, form an additional barrier that limits DNA accessibility to the transcriptional machineries. Moreover, by modulating StpA to H-NS ratios, bacteria might be able to dynamically fine-tune the selective permeability of this barrier and with that the recruitment of other regulatory proteins^75,86^. In conclusion, we demonstrate that controlling the ratio of nucleoid protein paralogs with distinct phase behaviors allows dynamic modulation of bacterial functional chromatin, thereby facilitating bacterial stress response and adaptation.

## LIMITATIONS OF THE STUDY

We present evidence for the key structural differences that contribute to the observed differences in H-NS and StpA’s phase behaviors. However, we acknowledge that the precise mechanism by which H-NS or H-NS/StpA fibrils phase separate is still unknown. Solving the high-resolution structures of the mixed H-NS/StpA fibrils would help in the understanding of the mechanism of phase separation. We also did not consider the possible involvement of other H-NS-interacting proteins (such as Hha) or posttranslational modifications in condensate stability under stress conditions which remains the topic for future studies.

## Acknowledgements

We thank Ken Wan for purifying the proteins and Lydia Freddolino for many helpful discussions. This work was supported by the NIH grant GM122506 to U.J., the NIH T32GM140223 to T.S.B., and the NIH DP2GM150019 to S.M. J.C.A.B and B.S. are supported by the Howard Hughes Medical Institute. Research reported in this publication was supported by the NIH S10OD030275 and the Arnold and Mabel Beckmann Foundation grants to the University of Michigan Cryo-EM Facility (U-M Cryo-EM). U-M Cryo-EM is grateful for support from the U-M Life Sciences Institute and the U-M Biosciences Initiative.

## Author Contributions

**J.G.:** Conceptualization, Methodology, Writing-Original draft preparation; Validation; Formal analysis, Investigation, Visualization, Supervision; **B.S.:** Investigation, Visualization,; **T.B.** Investigation, Visualization; Formal analysis; **P.M.:** Investigation, Visualization**; A.B.:** Investigation; **S.M.:** Supervision, Project administration, Funding acquisition; **J.B.** Supervision, Funding acquisition, Editing; **U.J.** Conceptualization, Supervision, Reviewing and Editing, Project administration, Funding acquisition.

## STAR METHODS

### Media and culture conditions

Bacteria were growth in either Luria-Bertani (LB) broth (rich media) or Gutnick broth (33.8 mM KH_2_PO_4_, 77.5 mM K_2_HPO_4_, 5.74 mM K_2_SO_4_, 0.41 mM MgSO_4_) supplemented with 0.4% w/v glucose, 10 mM NH_4_Cl, 10 µM ferric citrate and MOPS micronutrients ^87^ (hereafter referred as Gutnick+ medium). Unless otherwise specified, the growth temperature was set to 37°C and shaking was set to 200 rpm. For imaging and sampling purposes, overnight cultures of bacteria were inoculated into fresh media to an OD_600_ of 0.003. Exponentially growing cells were imaged and/or collected when OD_600_ reached 0.2 (exponential growth) or 24h thereafter (late stationary phase)^14^.

### Bacterial strain construction

Bacterial strains and plasmid vectors used in this study are listed in Table S1. DNA amplifications were performed using Platinum Superfi II Polymerase (Thermo Fisher Scientific). Primers used in this study can be found in Table S2. Endogenous fluorescent protein fusion strains were constructed using λ-Red recombination^88^. Briefly, linear DNA fragments flanked by at least 50 bp homologous sequences were electroporated into competent *E. coli* str. K-12 substr. MG1655 carrying the pKD46 plasmid. Recombination was selected by kanamycin resistance. Positive colonies were confirmed by PCR screening and Sanger sequencing (Eurofins Genomics). Successfully constructed strains express the protein of interest fused to a fluorescent protein at the C-terminus and linked by a 14-aa peptide^89^.

Gene knockouts were performed using P1 transduction with donor strains from the Keio collection^90^, and positive colonies were validated by PCR screening. For removal of selection markers, pCP20 plasmids were electroporated into competent bacteria and expression was induced at 43°C^88^. Kanamycin-sensitive cells were selected for further experiments. To monitor H-NS mediated gene repression, selected H-NS-silenced genes^46^ were C-terminally tagged with 3xFLAG peptide at their endogenous loci. For quantification of gene repression, FLAG-tagged genes were P1 transduced into various *hns*/*stpA* mutants.

### Growth assays

Overnight bacterial cultures were inoculated into fresh Gutnick+ medium and grown at 37°C with shaking at 200 rpm. At mid-logarithmic phase, cultures were back-diluted to the same initial OD_600_ and loaded into a transparent 96-well plate (Corning Costar) at a volume of 150 µl per well. Growth curves under the indicated conditions were generated by measuring OD_600_ every 10 minutes on a TECAN Spark® plate reader at the indicated temperatures using orbital shaking. Gutnick+ medium containing no bacteria was used for normalization.

### Live cell imaging and *in vivo* FRAP measurements

Bacterial cultures were loaded onto a glass coverslip and fixed with an agarose pad. Nucleoids were stained with 1 µg/ml 4′,6- diamidino-2-phenylindole (DAPI) for 15 minutes. Fluorescence and differential interference contrast (DIC) imaging was performed on a Leica SP8 inverted confocal microscope controlled by Leica LAS X software with white light laser, 405 nm diode laser, PMT detector, HyD detector and 100x oil objective lens. DAPI was imaged using 405 nm excitation. mNeonGreen was imaged using 505 nm excitation. mCherry was imaged using 592 nm excitation. Kymographs showing H-NS/StpA colocalization during exponential growth were based on images acquired from an MI-SIM super resolution microscope (CSR Biotech), using the DAPI, GFP and RFP channels. Intensity plots were generated using Fiji and Graphpad Prism. *In vivo* FRAP was performed with LAS X’s Zoom In mode. Circular regions of interest with diameter = 0.5 µm were photobleached with 505 nm (mNeonGreen) or 592 nm (mCherry) lasers, respectively. Three images were captured prior to photobleaching to obtain the initial fluorescence intensity, and cells were imaged every five seconds for two minutes to track fluorescence recovery.

Fluorescence intensity change over time was background corrected, and data were fitted using Graphpad Prism to an exponential curve F=A(1-e^-t/τ^) to obtain the apparent recovery time τ.

### Immunoblotting

For the quantification of steady-state protein abundance *in vivo*, cultures at the indicated growth conditions were collected, washed and resuspended in SDS-polyacrylamide gel loading buffer (6.5 mM Tris-HCl pH 7.0, 10% v/v glycerol, 2% w/v SDS, 0.05% w/v bromophenol blue, 2.5% v/v β-mercaptoethanol). Samples were incubated at 95°C for 10 minutes and loaded onto a 4-12% NuPAGE Bis-Tris gel (Invitrogen). After running at 175V for 45 minutes, the gel was transferred to a polyvinylidene difluoride membrane (Bio-Rad) and blocked with EveryBlot blocking buffer (Bio-Rad) for 5 minutes with agitation. The membrane was then incubated at RT in blocking buffer containing 1:1000 diluted mouse anti-FLAG (Sigma), mouse anti-mNeonGreen (Proteintech) or rat anti-mCherry (Thermo Fisher Scientific) monoclonal antibody for one hour. After five washes with TBST (20 mM Tris-HCl pH 7.4, 150 mM NaCl, 0.1% v/v Tween-20), the membrane was incubated at RT in blocking buffer containing 1:10,000 diluted anti-mouse or anti-rat secondary antibody labeled with IRDye 680RD or 800CW (LI-COR Biosciences) for one hour. Following multiple washes in TBST, the membrane was imaged using the LI-COR Odyssey CLx imager. Gel quantification was performed with ImageJ, and normalization was performed against total protein by either Coomassie staining of a parallel gel, or by Ponceau staining of the same blot prior to blocking. Three biological replicates were included in all experiments.

### Fractionation of soluble and insoluble proteins

Following previously published protocols ^91^, cultures were collected and resuspended in 50 µl chilled lysis buffer (10 mM potassium phosphate pH 6.5, 1 mM EDTA, 20% w/v sucrose, 1 mg/ml lysozyme, 50 U/ml benzonase). After incubating the samples for 30 minutes on ice, samples were frozen at - 80°C and then re-thawed. Upon addition of 360 µl chilled buffer A (10 mM potassium phosphate pH 6.5, 1 mM EDTA) and protease inhibitor (ApexBio), the samples were transferred into 2 ml tubes containing 200 µl 425-600 µm glass beads (Sigma-Aldrich). The tubes were subjected to 30- minute shaking at 1,400 rpm at 4°C. Afterwards, 200 µl of the lysate was collected and centrifuged at 16,000 rcf for 20 minutes. The supernatant containing all soluble proteins was collected and mixed with ¼ volume of 100% v/v trichloroacetic acid. After incubation of the samples for 10 minutes on ice, the precipitant was washed thrice with chilled acetone, heated at 37°C for less than one minute and resuspended in 100 µl SDS-polyacrylamide gel loading buffer. The pellet containing the insoluble proteins was washed with buffer A, followed by buffer B (buffer A containing 2% v/v Nonidet P-40) and buffer A, and resuspended in 100 µl SDS-polyacrylamide gel loading buffer. Samples were then incubated at 95°C and subjected to immunoblotting as described as above.

### Protein purification

*E. coli* strains BL21 (DE3) carrying a pET-21a-H-NS-His6-SUMO plasmid or the mutant variants were grown in LB-media (plus ampicillin) at 37°C until OD_600_ of 0.5 was reached. To induce protein expression, 0.1 mM of IPTG was added to the culture. After overnight induction at room temperature, the cells were collected by centrifugation (20 min, 5,000xg, 4°C) and resuspended in lysis buffer, consisting of 40 mM Tris-HCl pH 8.0, 10 mM NaPO_4_ pH 8.0, 400 mM NaCl, 15 mM imidazole, 10% v/v glycerol 250 µM MgCl_2_, 0.5 µg/ml DNase I (Invitrogen), 0.5 µg/ml RNase A (Thermo Fisher Scientific), and cOmplete protease inhibitor (Roche). After sonication for seven minutes, the lysate was centrifuged twice for 30 minutes at 30,000xg. The supernatant was loaded onto an equilibrated HisTrap HP column (Cytiva). The column was washed with lysis buffer followed by lysis buffer containing 30 mM imidazole, and H-NS was eluted with lysis buffer containing 500 mM imidazole. The eluate was supplemented with ULP1 SUMO protease (Thermo Fisher Scientific) and dialyzed overnight against 40 mM Tris-HCl pH 8.0, 300 mM NaCl. The dialyzed sample was passed through an equilibrated HisTrap HP column again, and the flowthrough containing the cleaved protein was mixed 1:3 with 25 mM Tris-HCl pH 8.5 and loaded onto a HiTrap Q column (Cytiva). H-NS was eluted with a gradient from 9 to 80% ÄKTA pure buffer B (1 M NaCl, 50 mM NaPO_4_ pH 8.0). The purified sample was dialyzed against 25 mM Tris-HCl pH 7.5 and 300 mM NaCl, concentrated and stored at -80°C. For purification of StpA and StpA F21C, *E. coli* strains BL21 (DE3) carrying pET-21a-StpA or StpAF21C-GST were grown at 37°C in LB (plus ampicillin) until OD_600_ of 0.5 as reached. Then, 0.1 mM of IPTG was added and protein production was induced at room temperature for 4 hours. The cells were pelleted by centrifugation (20 min, 5,000xg, 4°C) and lysed by sonication for 8 minutes in buffer containing 40 mM Tris-HCl pH 8.0, 10 mM NaPO_4_ pH 8.0, 500 mM NaCl, 5 mM DTT, 10% v/v glycerol, 250 µM MgCl_2_, 0.5 µg/ml DNase I (Invitrogen), 0.5 µg/ml RNase A (Thermo Fisher Scientific) and cOmplete protease inhibitor (Roche). The lysates were centrifuged twice for 30 minutes at 37,500xg. The supernatant was loaded onto an equilibrated 5 ml GSTrap FF column (Cytiva). The column was washed with lysis buffer and StpA was eluted with 500 mM imidazole in lysis buffer. The eluate was digested with HRV 3C protease (Thermo Fisher Scientific) and dialyzed against 40 mM Tris-HCl pH 7.5 and 500 mM NaCl overnight at 4°C. The dialyzed sample was passed through an equilibrated 5 ml GSTrap FF column (Cytiva). The flowthrough was mixed 3:25 with 50 mM NaPO_4_ pH 6.0 and passed through a HiTrap SP HP column (Cytiva). StpA and StpA F21C were eluted with a gradient from 9 to 80% ÄKTA pure buffer B. The purified sample was dialyzed against 25 mM Tris-HCl pH 7.5 and 500 mM NaCl, concentrated and stored at -80°C.

### *In vitro* protein and DNA labeling

To fluorescently label the purified proteins, 1 mg of the purified proteins was reduced at room temperature for 10 minutes with 180 µg Tris-(2- carboxyethyl) phosphine (TCEP) hydrochloride (Thermo Fisher Scientific) in buffer containing 50 mM K_2_HPO_4_ and 100 mM (for H-NS) or 500 mM (for StpA-F21C) NaCl. A pack of Cy3 (for H-NS) or Cy5 (for StpA-F21C) Maleimide Mono-Reactive Dye (Amersham) dissolved in 50 µl DMSO was supplemented to the reduced protein solution to a final volume of 1 ml. The solution was incubated for two hours at room temperature followed by an overnight incubation at 4°C. For removal of any excess dye, PD-10 desalting columns packed with Sephadex G-25 resin (Cytiva) were used following the manufacturer’s instructions. The protein eluates were concentrated with Amicon Ultra Centrifugal Filters of 3 kDa molecular weight cutoff. The protein concentration and labeling efficiency were calculated using a molecular extinction coefficient of 150,000 M^-1^cm^-1^ at 552 nm for Cy3 and at 650 nm for Cy5, with a correction factor at 280 nm of 0.08 for Cy3 and 0.05 for Cy5 following manufacturer’s guidelines. To label DNA, primers with 5’-FAM labels (IDT) were used to amplify DNA fragments of defined lengths (150 bp, 400 bp, 1 Kbp, 2.5 Kbp) from the *waa* operon ^14^ by colony PCR. The product size was validated by agarose gel electrophoresis and labeled DNA products were purified by phenol-chloroform precipitation.

### *In vitro* condensate formation and FRAP analysis

Purified H-NS and/or StpA proteins were diluted to the indicated concentrations in 100 mM monosodium L-glutamate, 1 mM magnesium acetate and 20 mM Tris-acetate pH 7.5 (GMT-buffer). Due to the NaCl concentration present in the respective protein storage buffers, samples containing H-NS alone, StpA alone or both proteins contained 30, 50 or 80 mM NaCl, respectively. Additional NaCl was added to ensure equal osmolarity when necessary. To test condensate stability, NaCl was added at the indicated concentrations. Samples were prepared in CS16-CultureWell removeable chambered coverglass (Grace Bio-Labs) treated overnight with 5% w/v pluronic acid (Thermo Fisher Scientific) and washed extensively with ddH_2_O. Imaging was performed on the same SP8 confocal microscope as described above. FAM, Cy3 and Cy5 were visualized using 495, 532 and 647 nm lasers, respectively. Phase transition was visualized with timelapse imaging. FRAP samples were incubated at room temperature for 30 minutes prior to imaging. For photobleaching of entire condensates, droplets with a diameter of ∼ 5 µm were selected. For partial photobleaching, regions with a diameter of ∼ 0.5 µm were selected among droplets with diameter of > 10 µm. FRAP settings were the same as described above.

### Preparation of condensates for turbidity measurements

H-NS or StpA at the indicated concentrations were prepared at room temperature as described above and supplemented with the indicated concentrations of DNA and/or NaCl. 100 µl of each sample was loaded into a transparent 96-well plate (Corning Costar) and absorbance at 350 nm was measured with a TECAN Spark® plate reader. Buffer containing no protein was used for normalization. All measurements were conducted in triplicates.

### Gene repression studies

*E. coli* MG1655 encoding select 3xFLAG or mCherry tagged H-NS- silenced genes^46^ at their endogenous loci were grown in Gutnick+ medium at 37°C or 42°C to mid-exponential phase, collected by centrifugation at 4000g, and subjected to immunoblotting as described above.

### Cryo-electron tomography of condensates

#### Plunge-freezing condensates

200 mesh copper grids with R2/1 holey carbon film (Quantifoil GmbH) were glow discharged for 30 s at 5 mA on a Pelco EasiGlow system. 3 μL of either H-NS alone, H-NS+DNA, or H-NS+DNA+StpA condensate-containing mixture was applied to grids in the environmental chamber of a Vitrobot Mark IV (Thermo Fisher Scientific) maintained at 24 ^°^C and 90% humidity. The grids were back- side blotted for 2.5 s with a blot force of 0 and plunge-frozen in liquid ethane. Frozen grids were clipped into Autogrids (Thermo Fisher Scientific) and stored in liquid N_2_ until further use.

#### Tilt-series acquisition & reconstruction

Grids containing condensates were imaged using a Titan Krios G4i transmission electron microscope (Thermo Fisher Scientific) operated at 300 kV, equipped with a K3 direct electron detector (Gatan Inc.) and a Bioquantum imaging filter (Gatan Inc.). Tilt series were acquired by a dose-symmetric scheme^92^ from +51° and -51° starting at 0° with 3° increments. Tilt series were acquired at a nominal magnification of 53,000x, equating to a pixel size of 1.681 Å/pixel and a filter slit width of 20 eV, using SerialEM software^93^ and scripts from PACEtomo^94^. Total dose per tilt series was ∼160 e^-^/Å^2^. The tilt series were aligned using patch tracking in IMOD^95^. All other reconstruction steps were performed using WarpTools software (https://doi.org/10.5281/zenodo.13982246). Reconstructed tomograms were denoised using cryoCARE^96^.

#### Fibril parameters estimation

To calculate approximate fibril lengths, denoised tomograms were first converted to 8-bit type using the IMOD mrcbyte tool^95^. 8-bit tomograms were loaded into Fiji^97^, adjusted to remove outlier pixels using the ‘Remove Outliers’ feature of the Integrated Image Filters Fiji plugin^98^, adjusted for enhanced contrast, and thresholded to achieve binarized tomograms. Individual central slices were extracted from representative tomograms and skeletonized using the Fiji plugin Skeletonize3D^99^. Longest shortest paths were calculated from skeletonized tomogram slices using the AnalyzeSkeleton Fiji plugin^100^. Fibril widths were measured manually using the measure feature in 3dmod from the IMOD package^95^. 101 measurements were performed per sample/condition. Fibril width and Longest Shortest Path jitter plots were generated using the ggplot2 system within RStudio (version 2025.5.1.513).

#### Negative stain transmission electron microscopy

Samples were stained with 0.75% (w/v) uranyl formate onto formavar coated 400 mesh copper grids stabilized with evaporated carbon film (Electron Microscopy Sciences). Briefly, 5 μL of sample was loaded onto glow discharged grids and were wicked away with Whatman paper after 45 seconds. Grids were washed twice with ddH_2_O and once with uranyl formate solution before incubating in uranyl formate for 45 seconds. Excess stain was wicked away with Whatman paper ad grids were imaged at room temperature using Fei Technai T12 microscope operating at 120kV. Images were captured at 50,000x with a sampling of 2.1 Å/pixel.

#### Molecular dynamics (MD) simulations of H-NS and StpA assemblies

Structural models of dimeric and tetrameric assemblies of *E. coli* H-NS and StpA proteins were generated using the Chai Discovery platform and the AlphaFold server^55^. The linear end-to-end truncated dimers were constructed from residues 1-82 of H-NS and 1-83 of StpA, forming the canonical dimerization site 1 and site 2 interfaces as reported previously^48,57,58^. Higher-order oligomers, including the homo-tetramers of H-NS (residues 1-137) and StpA (residues 1-134), as well as a hetero-tetrameric complex comprising two subunits of H-NS and two of StpA, were built using the same approaches. All initial models adopted extended conformations consistent with experimentally inferred linear arrangements of H-NS and StpA oligomerization domains. All-atom MD simulations were prepared with CHARMM-GUI^101^ using the CHARMM36m force field^102^ and TIP3P water model. A rectangular box of ∼25 × 15 × 15 nm was used for tetrameric complexes, and ∼13 × 13 × 13 nm was used for the truncated dimers. Each system was neutralized with 0.5 M KCl, a concentration chosen to probe the effect of elevated salt on complex stability. MD simulations were carried out using GROMACS version 2025.2^103^. After steepest-descent energy minimization, restrained equilibration was performed for 2.5 ns under constant particle number, pressure, and temperature (NPT ensemble). Production simulations were run for 200 ns at 310.15 K with a 2-fs integration time step and periodic boundary conditions. Temperature was controlled using the Nosé-Hoover thermostat, and pressure was maintained with the Parrinello-Rahman barostat. Long-range electrostatics were treated with the particle mesh Ewald (PME) method, with a real-space cutoff of 1.2 nm. Van der Waals interactions were smoothly switched off between 1.0 and 1.2 nm. All covalent bonds involving hydrogens were constrained with the linear constraint solver (LINCS) algorithm^104^. Trajectory analyses were performed with GROMACS utilities, including root mean square deviation (RMSD) and root mean square fluctuation (RMSF). Structural ensembles and conformational transitions were further examined using UCSF ChimeraX^105^ and Discovery Studio Visualizer studio^106^. Twenty snapshots from each MD trajectory were extracted at 10 ns intervals for structural comparison, and representative final MD structures at 200 ns were selected for figure preparation.

---

**Figure S1.**
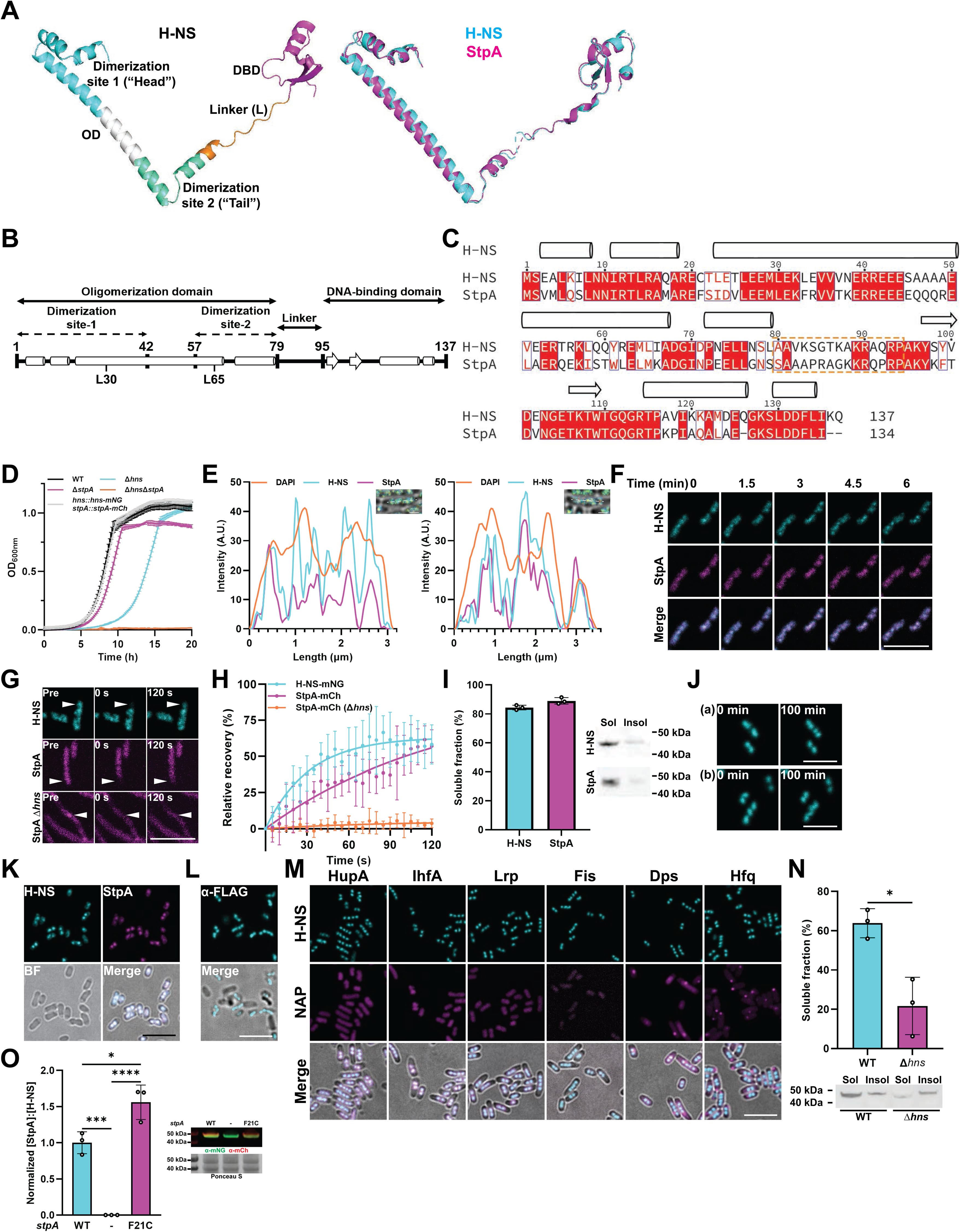
*In vivo* phase behavior of the *E. coli* paralogs H-NS and StpA (related to Figure 1). **A. Left:** Three-dimensional structure of *E. coli* H-NS as predicted by AlphaFold, with different domains highlighted. **Right:** Structure alignment of H-NS (cyan) and StpA (magenta) based on AlphaFold3 predicted structures and performed using jFATCAT (flexible). **B.** Domain organization and functional delineation of H-NS and StpA. **C.** Sequence alignment of *E. coli* H-NS and StpA. Red boxes with white letters indicate identical residues, white boxes with red letters indicate residues with similar properties. **D.** Growth curves of *E. coli* MG1655 WT, Δ*hns*, Δ*stpA* or Δ*hns/stpA* or MG1655 co-expressing H-NS-mNG and StpA-mCh from their respective endogenous loci, at 37°C in Gutnick+ minimal media. **E.** Representative kymographs of two bacteria at exponential phase, showing fluorescence intensity of DAPI (DNA), mNG (H-NS) and mCh (StpA) across the long axis of each cell. Scale bar is 1 µm. **F.** Live cell time course images of exponential H-NS-mNG and StpA-mCh expressing bacteria grown in Gutnick+ minimal media at 37°C. **G.** FRAP images of H-NS-mNG or StpA-mCh microdomains in exponentially grown WT *E. coli* (upper and middle panel) or StpA-mCh in the Δ*hns* strain (lowest panel). Frames depict puncta before (pre), immediately after (0s) and 120 sec after photobleaching. Arrowheads depict the photobleached areas. **H.** Quantification of H-NS-mNG (cyan) and StpA-mCh (magenta) FRAP signals of bacteria shown in (F). Mean, SD, *n* = 4-5. **I.** Solubility fractionation of endogenously expressed H-NS-mNG and StpA-mCh in stationary phase *E. coli*. A representative western blot image depicting soluble (Sol) and insoluble (Insol) fractions is shown. **J.** Live cell fluorescence time course images of H-NS-mNG in *E. coli* grown to stationary phase in Gutnick minimal media at 37°C. **K.** Live cell fluorescence and BF images of H-NS-mNG and StpA-mCh expressing *E. coli* grown to stationary phase in LB media at 37°C. **L.** Immunofluorescence and merged BF images of stationary phase H-NS-3xFLAG expressing *E. coli*. Cells were fixed and immunofluorescent detection was conducted using anti-FLAG antibodies. **M**. Fluorescence and merged BF images of stationary phase *E. coli* expressing H-NS-mNG (cyan) and select mCh-tagged *E. coli* NAPs (magenta) including HupA-mCh, IhfA-mCh, Lrp-mCh, Fis-mCh, Dps-mCh and Hfq-mCh from their respective endogenous loci. Lower panel depicts the merged overlay of mNG, mCh and BF channels. **N.** Solubility fractionation of endogenously expressed StpA^F21C^-mCh in exponential phase *E. coli*. A representative western blot image depicting soluble (Sol) and insoluble (Insol) fractions is shown. **O.** Normalized StpA:H-NS ratios in WT, Δ*stpA* and stpA^F21C^ strains. Quantification was performed by western blotting against mNG (H-NS) and mCh (StpA). Ponceau S stain against total protein was used for normalization. One-way ANOVA, mean, SD, *n* = 3, * *p* < 0.05, *** *p* < 0.001, **** *p* < 0.0001. A representative western blot image was shown. All scale bars 5 µm.

**Figure S2.**
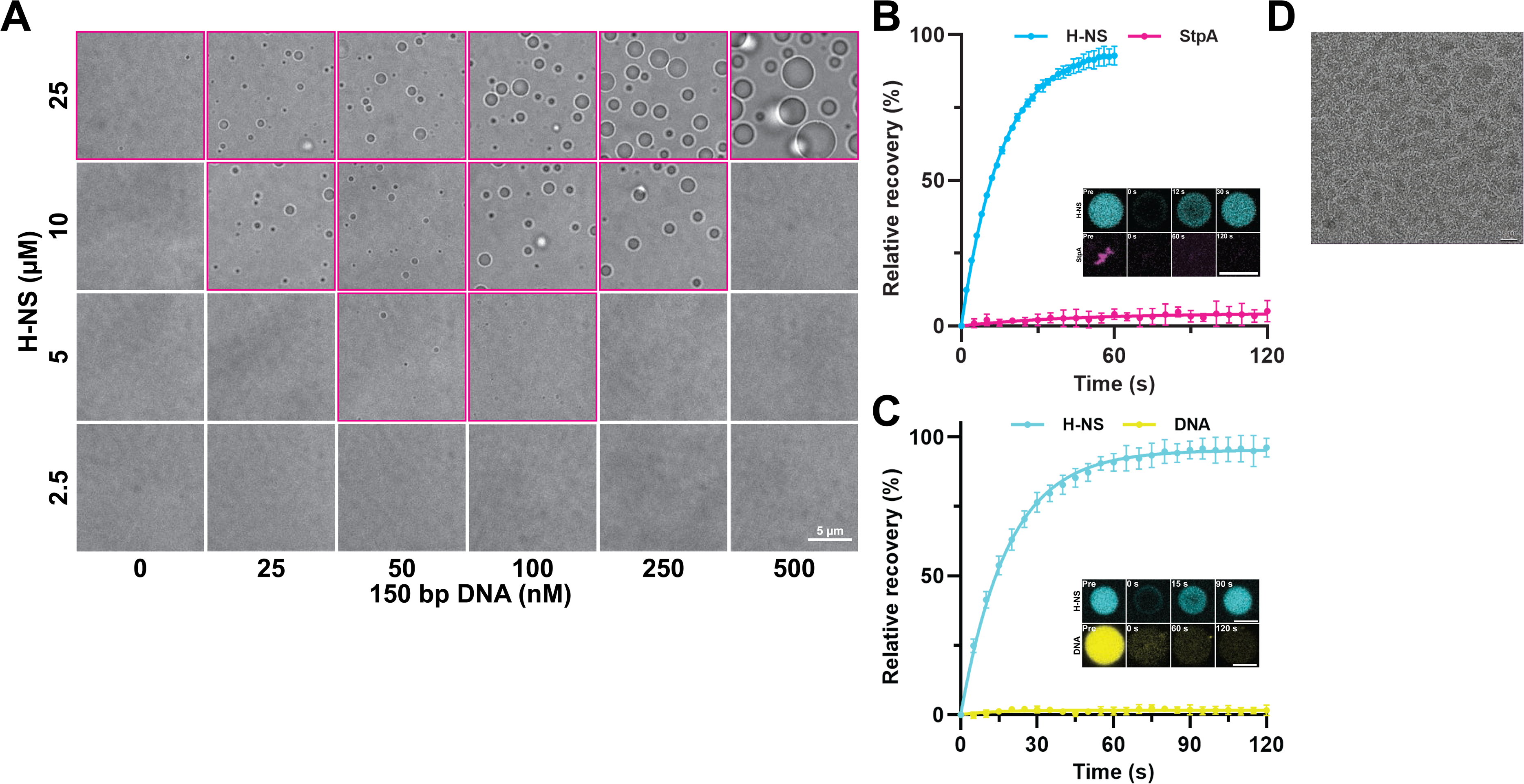
H-NS and StpA form co-condensates (related to Figure 2). **A.** BF images of droplets formed with various concentrations of H-NS and 150 bp DNA in GMT buffer to generate phase diagram shown in Fig. 2C. Panels containing droplets are highlighted in magenta. **B.** FRAP quantification of 100 µM H-NS (+ 1% Cy3-labeled) droplets or 100 µM StpA (+1 % Cy5-labeled StpA F21C) precipitates after bleaching the entire droplet (i.e., full FRAP); mean, SD, *n* = 4. **Inset:** Representative images of H-NS (above) or StpA (below), depicting before, immediately upon and indicated time points after photobleaching. **C.** Full FRAP quantification of 25 µM H-NS (+ 4% Cy3-labeled) mixed with 100 nM DNA (FAM-labeled) in co-condensates; mean, SD, *n* = 4. **Inset:** Representative images of H-NS and DNA depicting before, immediately upon and indicated time points after photobleaching. **D.** TEM analysis of 2.5 µM purified H-NS prepared in GMT buffer. Scale bar: 50 nm. Scale bars: 5 µm, unless otherwise indicated.

**Figure S3.**
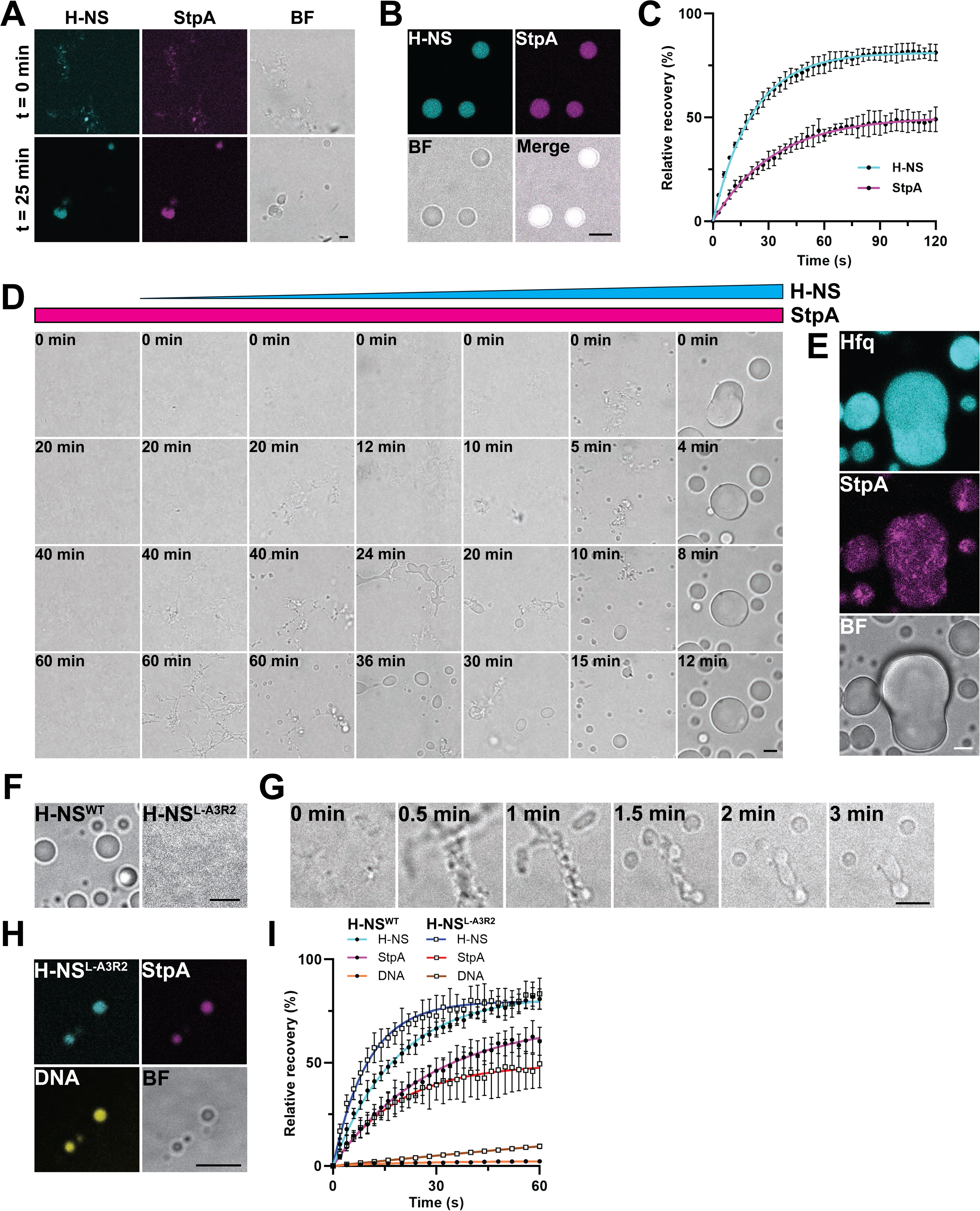
H-NS and StpA form co-condensates (related to Figure 3). **A.** Fluorescence and BF images of 25 µM H-NS (+ 4% Cy3-H-NS) and 10 µM preformed StpA (+ 4% Cy5-StpA^F21C^) fibrils immediately upon and 25 min after mixture in GMT buffer. **B.** Fluorescence images of the same samples as in (A) after 40 min of incubation. **C.** Quantification of partial FRAP of H-NS/StpA/DNA co-condensates shown in (B). Color code as before (mean, SD, *n* = 4-6). **Inset:** Representative images of FRAP time course. **D.** BF time course images of 10 µM StpA mixed with 0, 1, 2.5, 5, 10, 25 and 50 µM H-NS in GMT buffer. **E.** Fluorescence images of 100 µM Hfq (+ 1% Cy3-Hfq^S65C^) mixed with 10 µM preformed StpA fibrils in GMT buffer and incubated for 40 minutes. Scale bars: 5 µm. **F.** BF images of 25 µM H-NS or H-NS^L-A3R2^ mixed with 100 nM DNA in GMT buffer. **G.** BF time course images of 25 µM H-NS^L-A3R2^ and 100 nM DNA mixed with 10 µM pre-formed StpA fibrils in GMT buffer. **H.** Fluorescence and BF images of co-condensates containing 25 µM H-NS^L-A3R2^ (+ 4% Cy3-H-NS^L-A3R2^), 10 µM StpA (+4% Cy5-StpA^F21C^) and 100 nM FAM-DNA. **I.** Quantification of partial FRAP on co-condensates formed with 25 µM H-NS/H-NS^L-A3R2^ (+ 4% Cy3-H-NS/H-NS^L-A3R2^), 10 µM StpA (+ 4% Cy5-StpA^F21C^) and 100 nM FAM-DNA.

**Figure S4.**
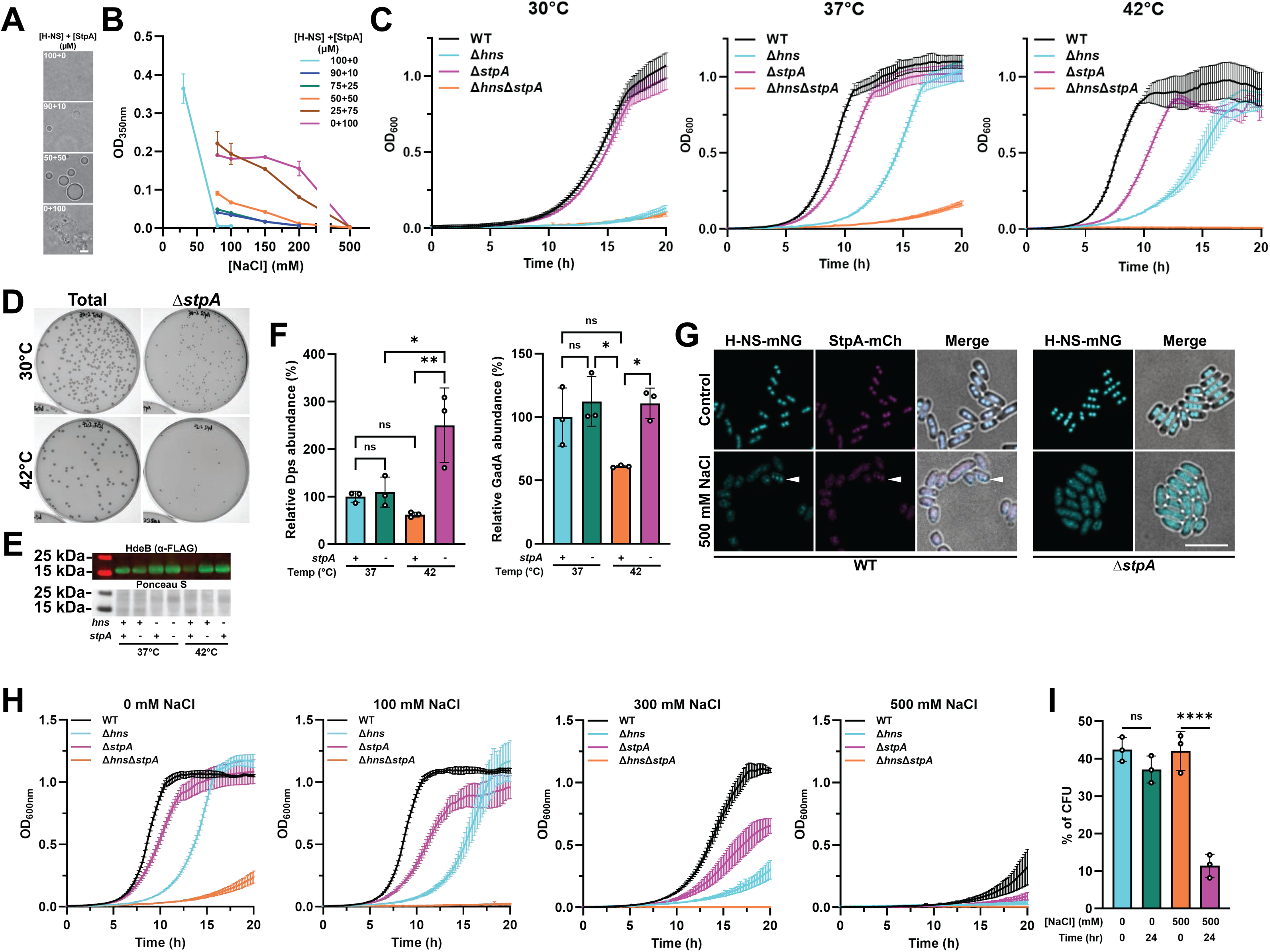
StpA is required for bacterial fitness under stress (related to Figure 4). **A.** BF images of H-NS and StpA mixed in GMT buffer at indicated concentrations in the presence of 100 mM NaCl. **B.** Turbidity of H-NS and StpA mixed at ths indicated concentrations in GMT buffer containing different amounts of NaCl. **C.** Growth curves of *E. coli* MG1655 WT, Δ*hns*, Δ*stpA* and Δ*hns*Δ*stpA* strains at the indicated temperatures. **D.** Representative images of *E. coli stpA::Kan* strain co-cultured with WT strain at 30°C or 42°C for 24h. Total CFU was estimated by plating serially diluted culture on LB agar, *stpA::Kan* CFU was estimated by plating on LB+Kan agar. Related to Fig. 4G. **E.** Representative western blot image related to Fig. 4H. **F.** Abundance of endogenously expressed Dps-mCh or GadA-3xFLAG in *E. coli* MG1655 WT or Δ*stpA* strains, grown to mid-exponential phase at 37°C or 42°C. Quantification was performed by western blotting against 3xFLAG (GadA) or mCh (Dps). Ponceau S staining of total protein was used for normalization; one-way ANOVA, *n* = 3, * *p* < 0.05, ** *p* < 0.01). **G.** Fluorescence and merged BF images of *E. coli* H-NS-mNG StpA-mCh or H-NS-mNG Δ*stpA* strains grown to stationary phase in Gutnick+ minimal medium containing 0 or 500 mM NaCl. Arrowhead highlights cells containing distinct foci. Scale bar: 5 µm. **H.** Growth curves of *E. coli* MG1655 WT, Δ*hns*, Δ*stpA* and Δ*hns*Δ*stpA* strains grown in Gutnick+ minimal medium containing indicated amounts of NaCl. **I.** Percentage of *stpA::Kan* strain CFU over total CFU before and 24h after co-culture with WT *E. coli* in Gutnick+ minimal medium containing indicated amounts of NaCl. Mean, SD, one-way ANOVA, *n* = 3, **** *p* < 0.0001.

**Figure S5.**
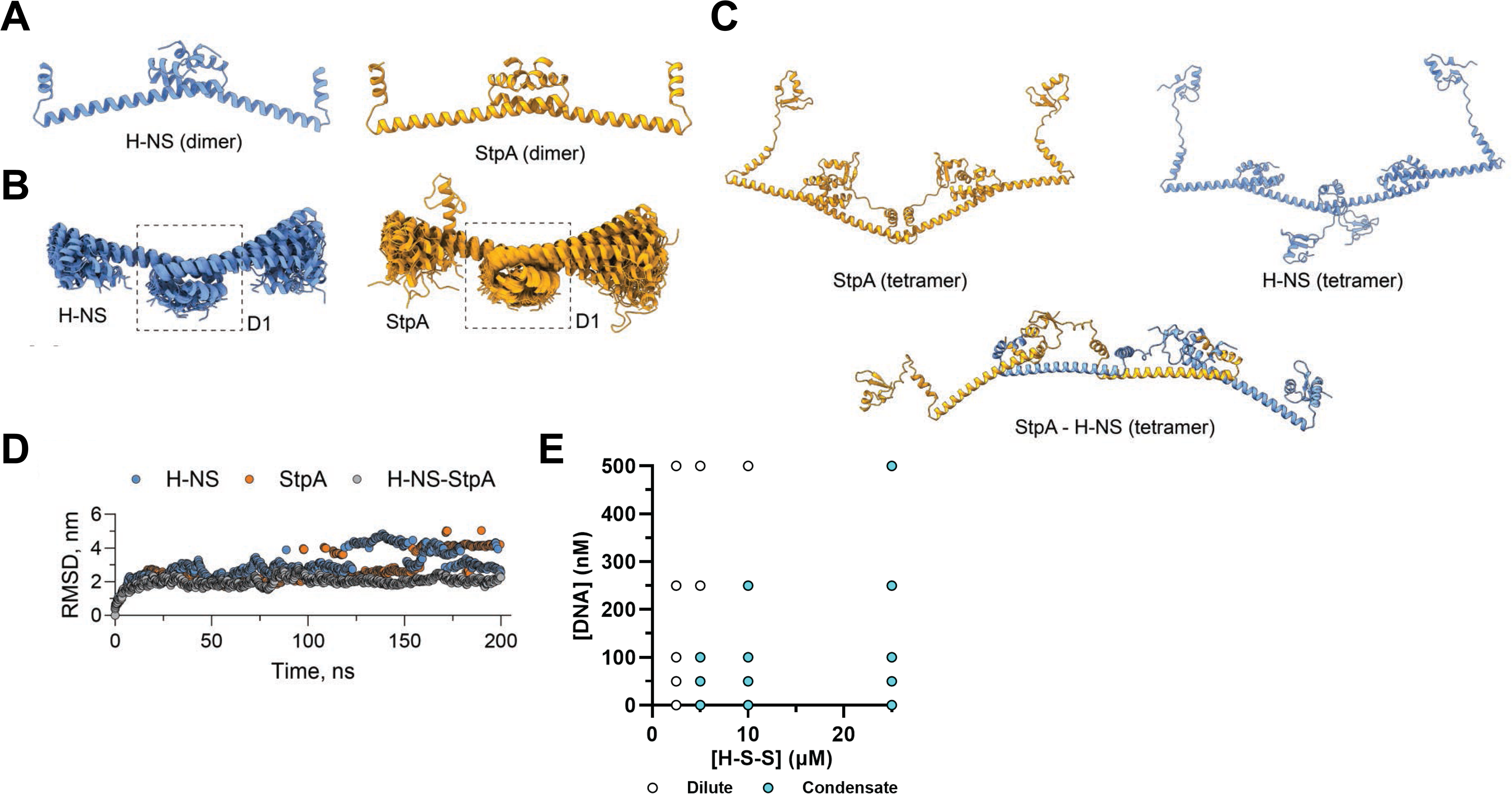
MD simulation of H-NS and StpA (related to Figure 5). **A.** Structure of N-terminally truncated H-NS (aa1-82) or StpA (aa1-83) dimers as predicted by AlphaFold. **B.** Ensembles of 20 superimposed snapshots (every 10 ns across a 200-ns run) for the N-terminally truncated dimers of H-NS and StpA at 500 mM KCl and 37°C. **C.** Structure of full-length H-NS homo-tetramer, StpA homo-tetramer or 2:2 H-NS:StpA hetero-tetramer as predicted by AlphaFold. **D.** Time evolution of root-mean-square deviation (RMSD) for full-length tetramers simulated under same conditions. **E.** Phase diagram of H-S-S and DNA in GMT buffer.

**Table S1.**
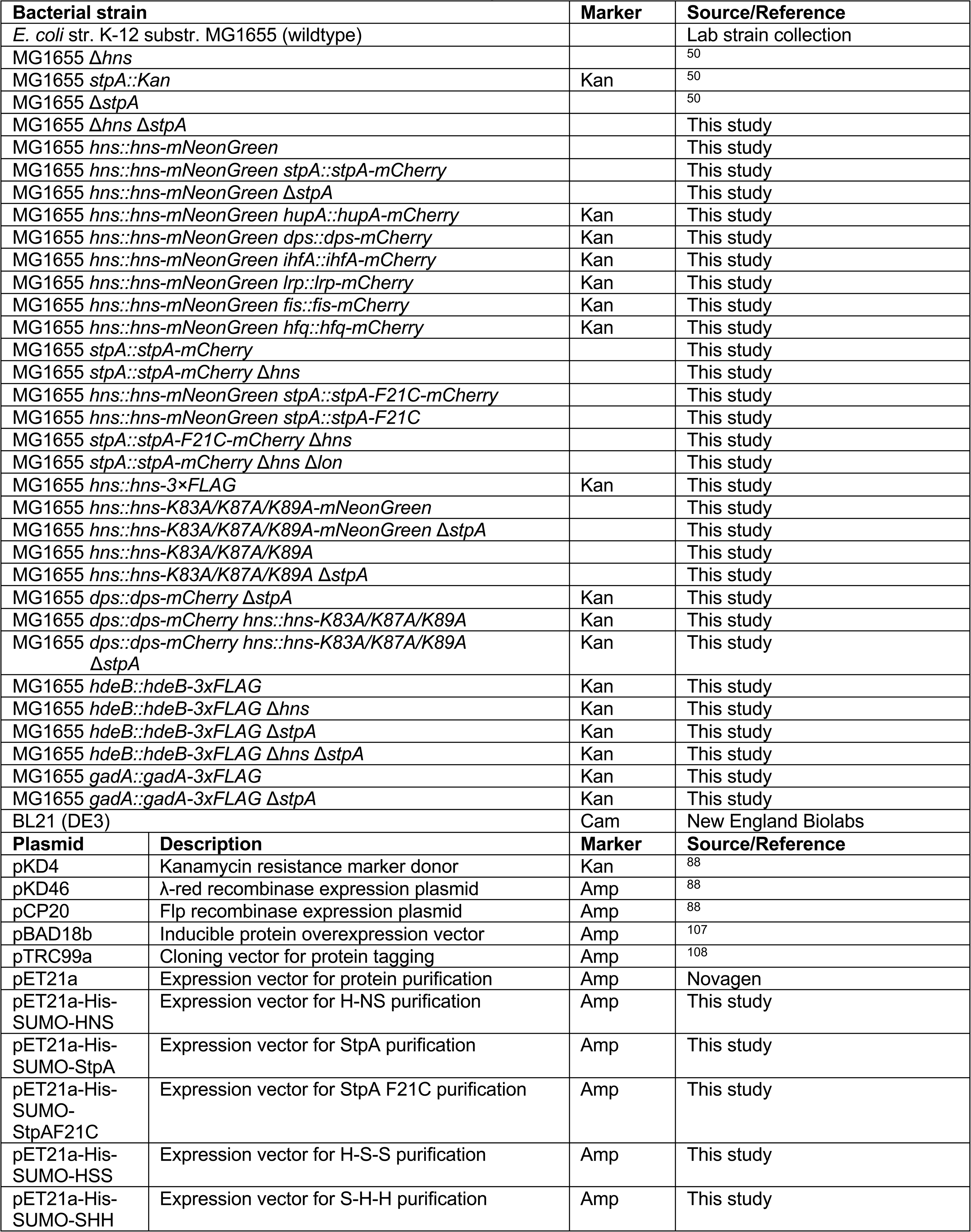
Strains and plasmid vectors used in this study.

| **Bacterial strain** |  | **Marker** | **Source/Reference** |
| --- | --- | --- | --- |
| *E. coli* str. K-12 substr. MG1655 (wildtype) |  |  | Lab strain collection |
| MG1655 $\Delta hns$ | | | 50 |
| MG1655 *stpA::Kan* |  | Kan | 50 |
| MG1655 $\Delta stpA$ | | | 50 |
| MG1655 $\Delta hns \Delta stpA$ | | | This study |
| MG1655 *hns::hns-mNeonGreen* |  |  | This study |
| MG1655 *hns::hns-mNeonGreen stpA::stpA-mCherry* |  |  | This study |
| MG1655 *hns::hns-mNeonGreen $\Delta stpA$* |  |  | This study |
| MG1655 *hns::hns-mNeonGreen hupA::hupA-mCherry* |  | Kan | This study |
| MG1655 *hns::hns-mNeonGreen dps::dps-mCherry* |  | Kan | This study |
| MG1655 *hns::hns-mNeonGreen ihfA::ihfA-mCherry* |  | Kan | This study |
| MG1655 *hns::hns-mNeonGreen lrp::lrp-mCherry* |  | Kan | This study |
| MG1655 *hns::hns-mNeonGreen fis::fis-mCherry* |  | Kan | This study |
| MG1655 *hns::hns-mNeonGreen hfq::hfq-mCherry* |  | Kan | This study |
| MG1655 *stpA::stpA-mCherry* |  |  | This study |
| MG1655 *stpA::stpA-mCherry $\Delta hns$* |  |  | This study |
| MG1655 *hns::hns-mNeonGreen stpA::stpA-F21C-mCherry* |  |  | This study |
| MG1655 *hns::hns-mNeonGreen stpA::stpA-F21C* |  |  | This study |
| MG1655 *stpA::stpA-F21C-mCherry $\Delta hns$* |  |  | This study |
| MG1655 *stpA::stpA-mCherry $\Delta hns \Delta lon$* |  |  | This study |
| MG1655 *hns::hns-3×FLAG* |  | Kan | This study |
| MG1655 *hns::hns-K83A/K87A/K89A-mNeonGreen* |  |  | This study |
| MG1655 *hns::hns-K83A/K87A/K89A-mNeonGreen $\Delta stpA$* |  |  | This study |
| MG1655 *hns::hns-K83A/K87A/K89A* |  |  | This study |
| MG1655 *hns::hns-K83A/K87A/K89A $\Delta stpA$* |  |  | This study |
| MG1655 *dps::dps-mCherry $\Delta stpA$* |  | Kan | This study |
| MG1655 *dps::dps-mCherry hns::hns-K83A/K87A/K89A* |  | Kan | This study |
| MG1655 *dps::dps-mCherry hns::hns-K83A/K87A/K89A $\Delta stpA$* |  | Kan | This study |
| MG1655 *hdeB::hdeB-3×FLAG* |  | Kan | This study |
| MG1655 *hdeB::hdeB-3×FLAG $\Delta hns$* |  | Kan | This study |
| MG1655 *hdeB::hdeB-3×FLAG $\Delta stpA$* |  | Kan | This study |
| MG1655 *hdeB::hdeB-3×FLAG $\Delta hns \Delta stpA$* |  | Kan | This study |
| MG1655 *gadA::gadA-3×FLAG* |  | Kan | This study |
| MG1655 *gadA::gadA-3×FLAG $\Delta stpA$* |  | Kan | This study |
| BL21 (DE3) |  | Cam | New England Biolabs |
| **Plasmid** | **Description** | **Marker** | **Source/Reference** |
| pKD4 | Kanamycin resistance marker donor | Kan | 88 |
| pKD46 | λ-red recombinase expression plasmid | Amp | 88 |
| pCP20 | Flp recombinase expression plasmid | Amp | 88 |
| pBAD18b | Inducible protein overexpression vector | Amp | 107 |
| pTRC99a | Cloning vector for protein tagging | Amp | 108 |
| pET21a | Expression vector for protein purification | Amp | Novagen |
| pET21a-His-SUMO-HNS | Expression vector for H-NS purification | Amp | This study |
| pET21a-His-SUMO-StpA | Expression vector for StpA purification | Amp | This study |
| pET21a-His-SUMO-StpAF21C | Expression vector for StpA F21C purification | Amp | This study |
| pET21a-His-SUMO-HSS | Expression vector for H-S-S purification | Amp | This study |
| pET21a-His-SUMO-SHH | Expression vector for S-H-H purification | Amp | This study |

**Table S2.**
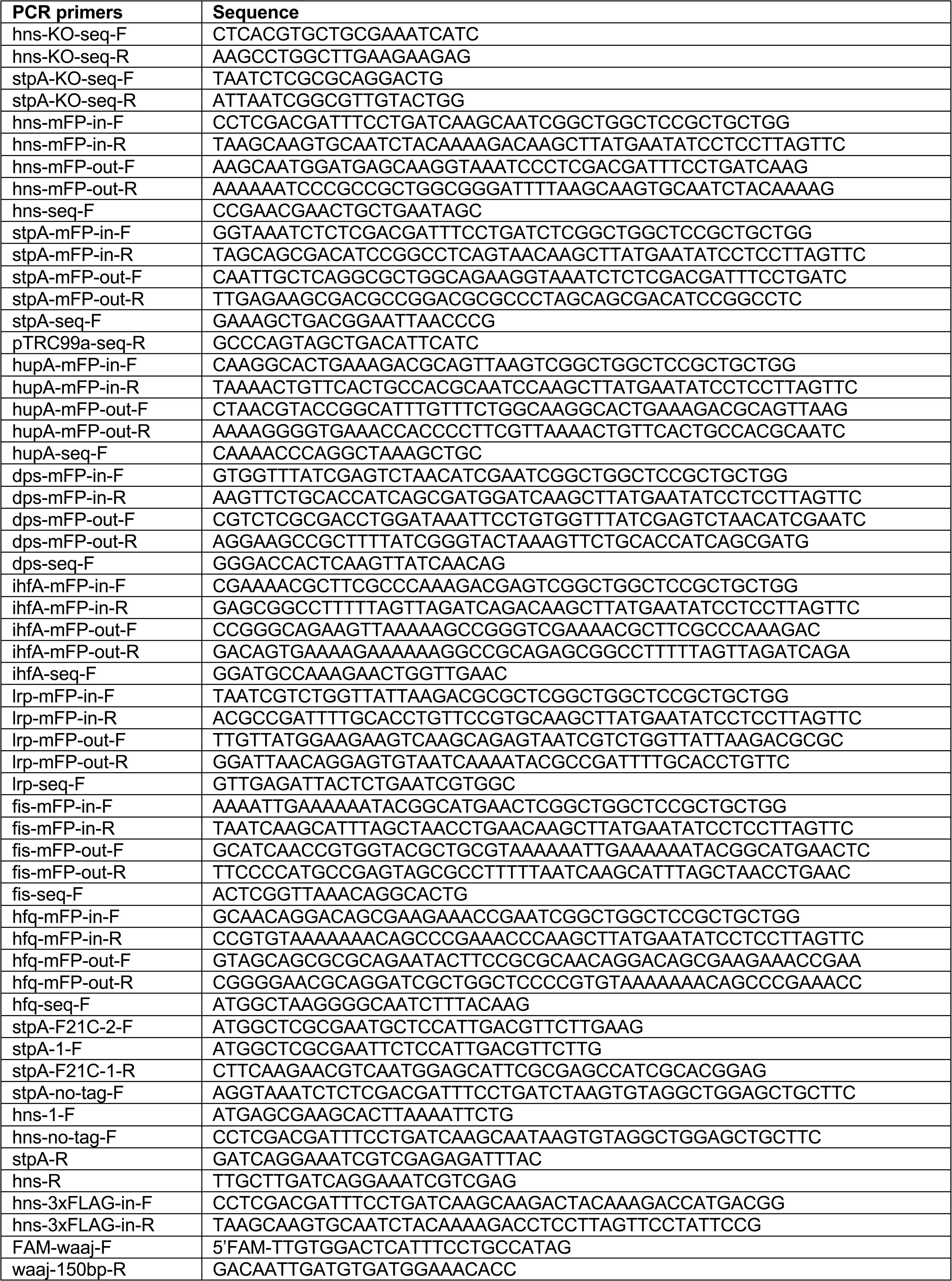
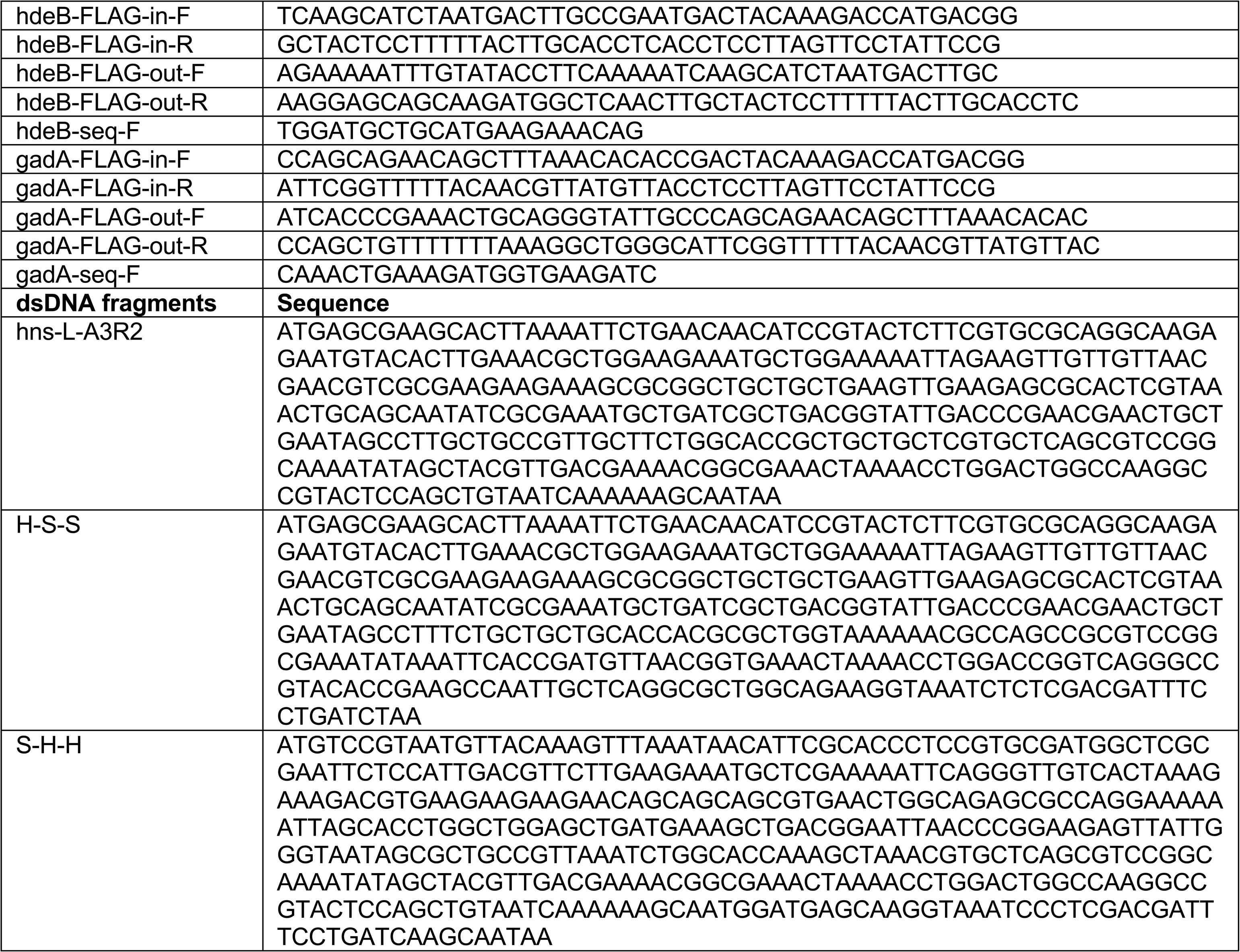
Primers used in this study.

